# Connection between drug-mediated neurotransmission and salivary microbiome

**DOI:** 10.1101/2023.08.20.553896

**Authors:** Daisuke Hisamatsu, Yusuke Ogata, Haruka Takeshige-Amano, Wataru Suda, Taku Hatano, Daisuke Asaoka, Yo Mabuchi, Yuna Naraoka, Nobuhiro Sato, Takashi Asada, Nobutaka Hattori, Masahira Hattori, Chihiro Akazawa

**Affiliations:** Intractable Disease Research Center, Juntendo University Graduate School of Medicine, 2-1-1 Hongo, Bunkyo-ku, Tokyo, 113-8421, Japan; Laboratory for Microbiome Sciences, RIKEN Center for Integrative Medical Sciences, Yokohama, Japan; Department of Neurology, Juntendo University Graduate School of Medicine, Tokyo, Japan; Department of Gastroenterology, Juntendo Tokyo Koto Geriatric Medical Center, Tokyo, Japan; Department of Microbiota Research, Juntendo University Graduate School of Medicine, Tokyo, Japan; Memory Clinic Ochanomizu, Tokyo, Japan

## Abstract

Salivary microbiome alterations associated with cognitive function have been reported in patients with neurodegenerative diseases^1–3^. Gut microorganisms can modulate therapeutic efficacy via drug metabolism^4–7^. Additionally, several drugs against diabetes and inflammatory bowel disease can lead to microbial dysbiosis^8–10^. However, the effect of central nervous system (CNS) drug use on the microbiome remains unknown. Here, we show that the usage of anti-dementia drugs, including donepezil and memantine, more largely affects the salivary microbiome composition than the gut microbiome composition. We observed salivary microbiome diversity reduction in patients with neurodegenerative diseases who received *N*-methyl-D-aspartate receptor inhibitor drugs. Furthermore, the use of acetylcholine-modulating drugs contributed to the salivary microbiome composition, suggesting that the salivary microbiome responds to changes in CNS drug-induced cerebral acetylcholine and glutamate levels. Multivariate analysis adjusted with or without the use of anti-dementia drugs demonstrated that the difference in the salivary microbiome correlated with cognitive function. We show the unique salivary microbiome structure of CNS drug users, suggesting the possibility of monitoring pharmacokinetics using the salivary microbiome. Our results also provide evidence of the presence of the microbiome–oral–brain axis and will accelerate the elucidation of the interplay between the salivary microbiome and neurodegeneration.

## Introduction

Mild cognitive impairment (MCI) is a preclinical stage of dementia, and half of the affected individuals transition to dementia, including Alzheimer’s disease^11,12^. Currently, 143 drugs are undergoing clinical trials for Alzheimer’s disease, most of which prevent or delay cognitive impairment^12^. However, to date, 99% of drug trials have failed^13^, partially due to inappropriate patient selection for individual therapies^14,15^. Dementia results from various primary diseases, including Alzheimer’s disease, dementia with Lewy bodies, and Parkinson’s disease^16^. Early diagnosis of the primary disease is essential for appropriate therapy as brain pathology progresses from the prodromal stage of the clinical symptoms in neurodegeneration^16,17^. Therefore, developing precise diagnostic modalities to provide new therapeutic technologies is essential. Saliva contains numerous diverse microorganisms^18^, constituting a salivary or oral microbiome associated with host physiologies, including systemic diseases and circadian rhythms^19–21^. In patients with neurodegenerative diseases, including Alzheimer’s disease and Parkinson’s disease, gut and salivary microbiomes shift in correlation with cognitive function^1–3,22–25^ or disease progression^26–29^. Recent meta-analyses at the population level have suggested that the use of therapeutic drugs induces gut microbiome alterations, particularly in gastroenterological and metabolic disorders^30–32^. Acetylcholinesterase inhibitors (AChEi) and *N*-methyl-D-aspartate receptor antagonists (NMDAra), such as donepezil and memantine, have been approved as anti-dementia drugs worldwide^33^. Anti-parkinsonian drugs also include amantadine HCl, an NMDAra, and anticholinergics that modulate the uptake of acetylcholine^34^. The use of anti-parkinsonian drugs, such as levodopa and dopamine agonists, stimulates the dopamine receptors to replace depleted dopamine, and some of these drugs are known to affect the microbiome in users^27^. Considering that central nervous system (CNS) drugs, including anti-parkinsonian and anti-dementia drugs, maintain brain physiology by regulating neurotransmitters^33,34^, we hypothesised that the use of anti-dementia drugs may affect the salivary microbiome in patients with neurodegenerative diseases. However, it is unclear whether AChEis and NMDAras affect gut and salivary microbiomes in neurodegeneration. Moreover, few studies have considered the effects of anti-dementia drug use on microbiomes and their correlation with cognitive function or disease progression. Finding a promising microbiome biomarker that correlates with cognitive function involves considering the influence of anti-dementia drugs, such as donepezil and memantine. To develop a new diagnostic biomarker based on the salivary microbiome profiles that enable the classification of neurodegenerative diseases, we demonstrated that CNS drugs for Alzheimer’s and Parkinson’s diseases affect the salivary and gut microbiomes.

## Results

### Altered salivary microbiome diversity

We initially performed bacterial 16S rRNA gene sequencing of saliva and stool samples from participants diagnosed with MCI and dementia using cognitive function tests. Here, we defined five metadata types as confounding factors for microbial variability: disease category, age, sex, cohort, and drug type (Fig. 1a). The participants were divided into two sets: a ‘whole’ set with a mixture of treated and untreated patients and an ‘untreated’ set with only untreated patients (Extended Data Fig. 1a,b and 2a,b). Diversity analyses showed an increase in α-diversity (Shannon index) in dementia compared to MCI in the whole set, although no significant change was observed in the untreated set (Extended Data Fig. 3a,b). In contrast, β-diversity (UniFrac distance) was significantly different between the healthy cognitive control (HC) and cognitive impairment groups, including the MCI and dementia groups (*P* < 0.05), but was not significantly different between the MCI and dementia groups in both the whole and untreated sets (whole set, *R*^2^ = 0.008, *P* = 0.062; untreated set, *R*^2^ = 0.012, *P* = 0.949; Extended Data Fig. 3c,d).

**Fig. 1.**
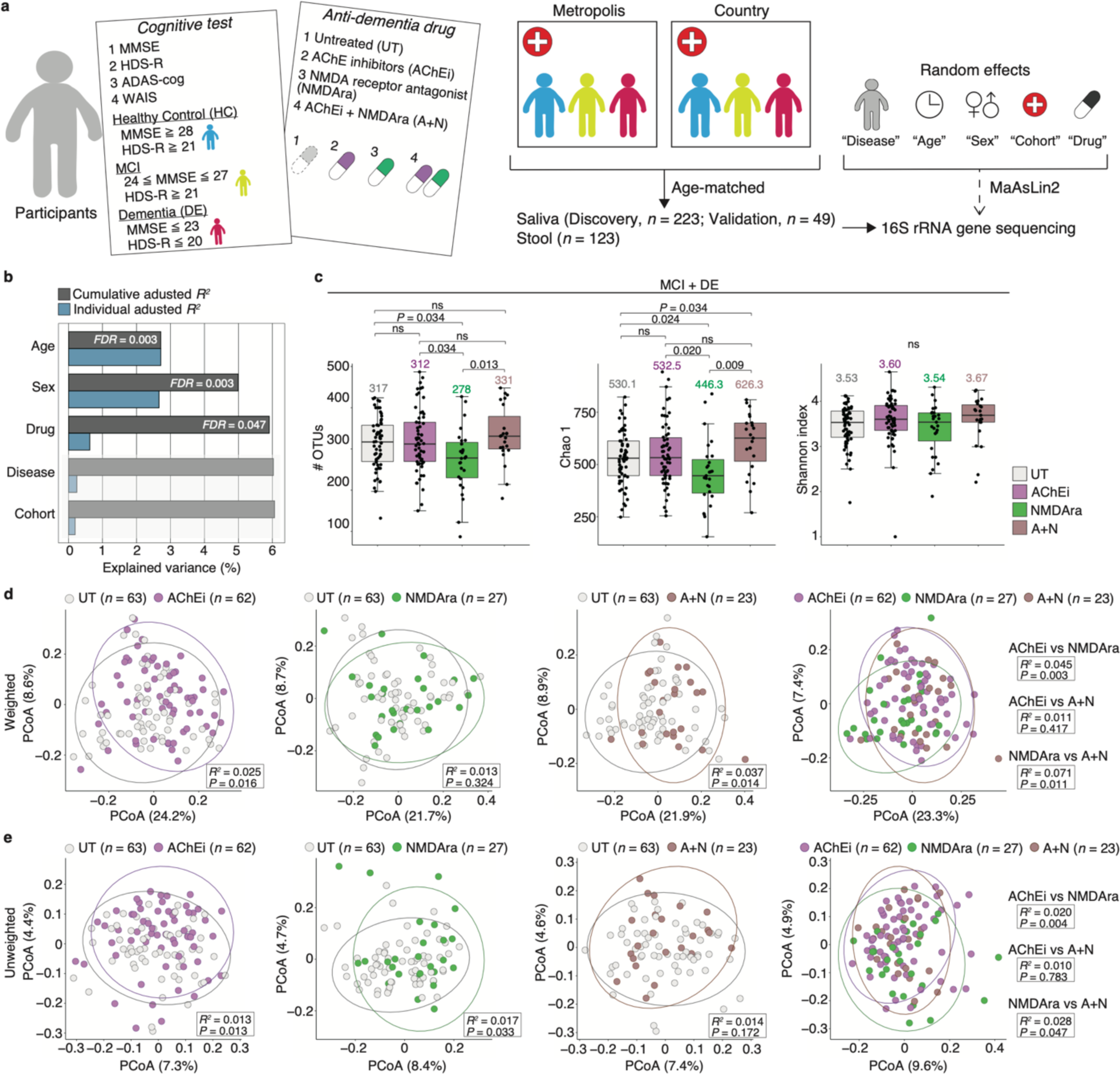
Contribution of anti-dementia drug use to the salivary microbiome diversity. **a,** Schematic diagram showing the study design. The primary cohort was recruited from two different regional hospitals (for the saliva analysis) and a metropolitan hospital (for the stool analysis). The saliva of the validation cohort was collected from a metropolitan hospital. Clinical data, including cognitive test scores and prescription drug information, were collected. **b,** Individual and cumulative adjusted *R*^2^ (explained variance) of covariates in stepwise RDA using weighted UniFrac distance in cognitive impairment groups (MCI and dementia; *n* = 175). Light colour indicates no significance (*FDR* > 0.05). **c,** Comparison of the observed OTU number and α-diversity score (Chao 1 and Shannon index) by drug category in cognitive impairment groups (MCI and dementia; *n* = 175). Each median is shown in the graph. Statistical significance was determined using the Wilcoxon rank-sum test with Benjamini–Hochberg correction (*P* < 0.05). ns, not significant. **d, e,** Weighted (d) and unweighted (e) UniFrac-PCoA between each drug category in the cognitive impairment group (MCI and dementia; *n* = 175). The number of participants in each drug category is depicted in the figure. The *R*^2^ and *P*-values were determined using permutational multivariate analysis of variance via the Benjamini–Hochberg method. Dots represent individual participants. MCI, mild cognitive impairment; OTU, operational taxonomic unit; PCoA, principal coordinate analysis; RDA, redundancy analysis.

The patients with MCI and dementia received uniform anti-dementia drugs, and therefore, the drug type rather than the disease category possibly determined the microbiome composition. Thus, we divided the cognitive impairment group (MCI and dementia) into four subgroups according to the type of anti-dementia drugs used^33^: untreated, AChEi-treated, NMDAra-treated, and AChEi plus NMDAra dual therapy (A+N) subgroups (Extended Data Fig. 1c). To assess the contributions of the five confounding factors to the microbiome composition, we performed stepwise redundancy analysis (RDA) based on the weighted UniFrac distance. Three variables significantly contributed to the overall microbiome composition in the cognitive impairment group (cumulative adjusted *R*^2^: age, 2.71%, *FDR* = 0.003; sex, 4.99%, *FDR* = 0.003; drug, 5.91%, *FDR* = 0.047; Fig. 1b), indicating that the alteration in the salivary microbiome was more largely influenced by the drug type than disease category (MCI or dementia). We then evaluated the α-diversity of the four subgroups and found a significant reduction in the number of observed operational taxonomic units (#OTUs) and Chao1 index in the NMDAra-treated subgroup compared with those in the untreated, AChEi-, and A+N-treated subgroups (#OTUs, *P* = 0.034, *P* = 0.034, *P* = 0.013; Chao1, *P* = 0.024, *P* = 0.020, *P* = 0.009; Fig. 1c). Furthermore, the β-diversity showed significant differences between the untreated and each treated subgroups (untreated vs AChEi-treated, *R*^2^ = 0.025, *P* = 0.016; untreated vs NMDAra-treated, *R*^2^ = 0.017, *P* = 0.033; untreated vs A+N-treated, *R*^2^ = 0.037, *P* = 0.014; Fig. 1d,e). Interestingly, despite the differences between two treated subgroups (AChEi vs NMDAra and NMDAra vs A+N), no significant differences were observed between the AChEi- and A+N-treated subgroups (Fig. 1d,e).

### Cognition-associated salivary microbiome

To evaluate the contribution of anti-dementia drugs to the cognitive function-related salivary microbiome, we performed a multivariate analysis adjusted with anti-dementia drug use on the salivary microbiome. We first compared the microbiome composition of individual patients in the disease category at the phylum, genus, and species levels (Fig. 2a and Extended Data Fig. 4a,b). The results found that several species significantly correlated with cognitive function in the untreated set, including those not observed in the whole set (Extended Data Fig. 4c,d,e). The drug type contributed significantly to microbiome composition at the genus level (cumulative adjusted *R*^2^ = 4.89%, *FDR* = 0.037; Fig. 2b). We also observed several genera of which the abundance differed by disease category in the untreated and whole sets (Fig. 2c,d,e). Interestingly, most of the genera significantly increased in abundance in the dementia group compared with the MCI group in the whole set but not in the untreated set in which, in contrast, some genera exhibited a decrease in abundance in the dementia group. Similar results were further observed for the correlation between the microbiome and cognitive function differed depending on the analysis with or without drug use (Fig. 2f). Furthermore, we assessed the correlation between the abundance of salivary species and Mini-Mental State Examination (MMSE) score, a conventional cognitive function test, in the untreated and whole sets. Spearman’s correlation coefficient analysis showed that the abundance of 11 species was significantly correlated with the MMSE score in the untreated set, although 7 of them were not found in the whole set (*P* > 0.05; Extended Data Fig. 5). Thus, multivariate analysis, with anti-dementia drug use as a confounding factor, may be essential in identifying biomarkers in the salivary microbiome to distinguish cognitive function.

**Fig. 2.**
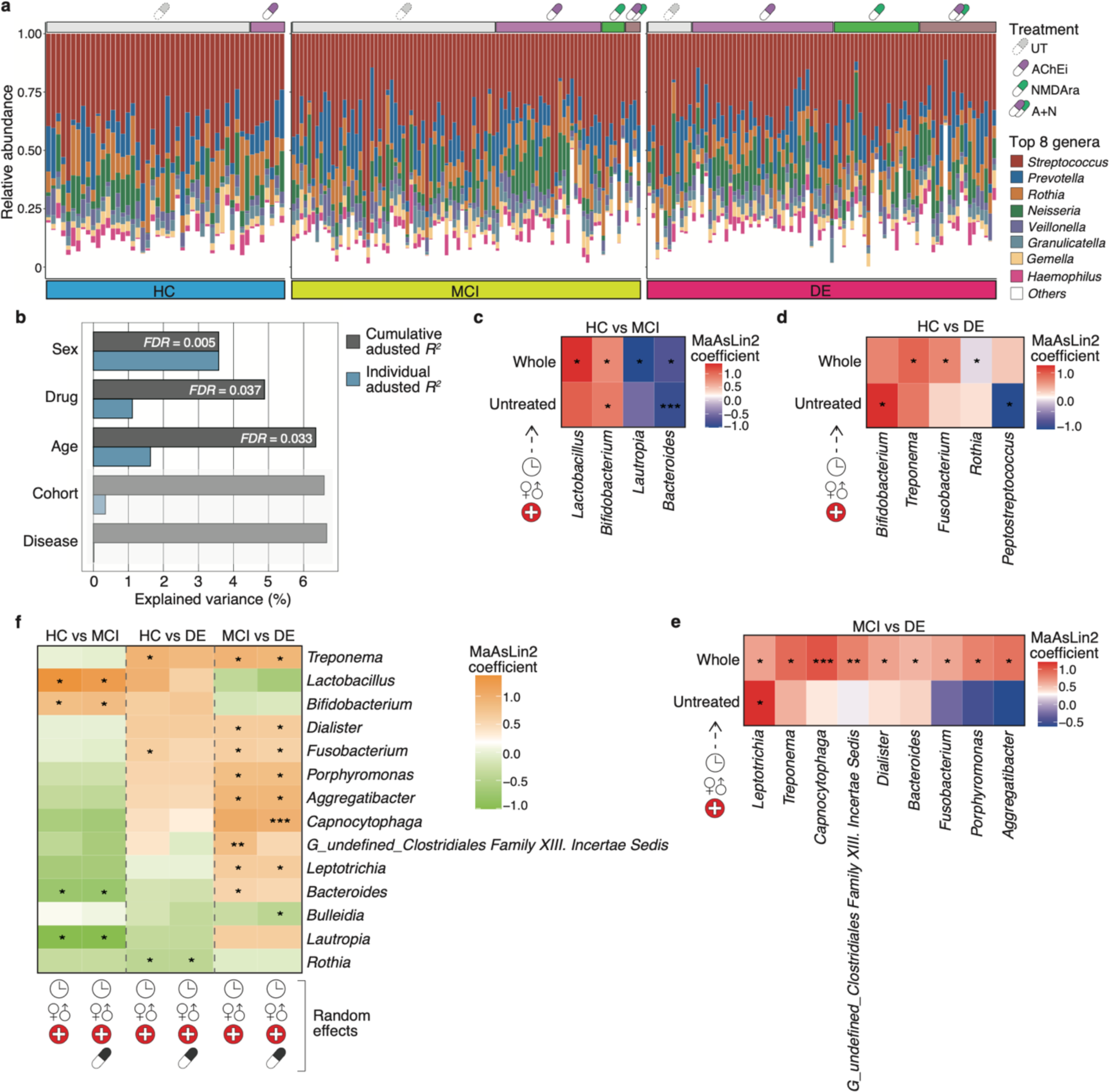
Considering anti-dementia drug use provides the precise microbiome composition associated with cognitive function. **a,** Relative abundance plots of salivary microbial genera across each disease category with drug category information (HC, *n* = 48; MCI, *n* = 89; DE, *n* = 86). Bars represent individual participants, and their colours show the top eight genera with the highest abundance of the 43 genera with a relative mean abundance > 0.1%. **b,** Individual and cumulative adjusted *R*^2^ (explained variance) of covariates in stepwise RDA using the genera composition of the salivary microbiome in cognitive impairment groups (MCI and dementia; *n* = 175). Light colour indicates no significance (*FDR* > 0.05). **c, d, e,** Heatmaps show representative genera significantly enriched and depleted between HC vs MCI (c), HC vs DE (d), and MCI vs DE groups (e) in whole (treated and untreated patients) and untreated sets (whole: HC, *n* = 48; MCI, *n* = 89; DE, *n* = 86; untreated: HC, *n* = 41; MCI, *n* = 52; DE, *n* = 11). Statistical significance was determined using the MaAsLin2 package with age, sex, and cohort as random effects (*P* < 0.05). **f,** Heatmap shows representative genera significantly enriched and depleted among each disease category in the whole set (HC, *n* = 48; MCI, *n* = 89; DE, *n* = 86). Statistical significance was determined using the MaAsLin2 package with age, sex, cohort, and/or drug as random effects (*P* < 0.05). \**P* < 0.05; \*\**P* < 0.01; \*\*\**P* < 0.005. HC, healthy control; MCI; mild cognitive impairment; DE, dementia.

Subsequently, we performed a random forest classification in both the untreated and whole sets using the same salivary species sets to examine the effect of anti-dementia drugs on the classification of neurodegenerative diseases. When the model was used to distinguish between the MCI and dementia groups with the highest predictive value in the untreated set, the area under the curve (AUC) showed a 0.82-fold reduction in the whole set (untreated set, AUC 0.823; whole set, AUC 0.675; Extended Data Fig. 6a). Moreover, the AUC tended to be higher in the NMDAra-treated subgroup than in the other three subgroups (untreated, AChEi-treated, and A+N-treated subgroups) in the cognitive impairment group (Extended Data Fig. 6b). Variable importance was observed for the different species in each set (Extended Data Fig. 6c,d,e). We demonstrated that the overall salivary microbiome composition was similar between patients with MCI and dementia (Extended Data Fig. 3c,d). Altogether, considering multiple confounding factors, including anti-dementia drug use, provided a precise classification of the cognitive impairment group by the salivary microbiome.

### Less influenced gut microbiome

To explore the effects of anti-dementia drug use on the gut microbiome, we first analysed α-diversity indices and found that they tended to increase in patients with MCI and dementia compared with HCs (Extended Data Fig. 7a,b). The β-diversity showed no significant change in the disease category (Extended Data Fig. 7c,d). In the cognitive impairment group, we performed stepwise RDA based on the weighted UniFrac distance, which demonstrated that sex was the only confounding factor among the four variables (sex, age, disease, and drug use) for the gut microbiome (Extended Data Fig. 8a). Although AChEi and NMDAra drugs are absorbed in the gut^35,36^, α-diversity and β-diversity exhibited no significant changes in each drug category of the cognitive impairment group (Extended Data Fig. 8b,c,d).

We next observed the gut microbiome composition at phylum, genus, and species levels in each disease category (Extended Data Fig. 9a,b,c). To evaluate the effect of anti-dementia drug use on cognitive function-correlated microbiome, we performed a multivariate analysis with or without considering anti-dementia drug use. We found that several bacteria correlated with cognitive function at the genus and species levels; however, most of these species were shared by both analyses with or without consideration of drug use as a confounder (Extended Data Fig. 9d,e), which was in contrast to the findings in the salivary microbiome (Fig. 2f). These results indicate less influence of anti-dementia drugs on the gut microbiome than on the salivary microbiome.

### Anti-dementia drug user-specific microbiome

To verify the effects of anti-dementia drug use on the salivary microbiome of the primary cohort, we further investigated saliva samples from patients with MCI and dementia in a validation cohort (Extended Data Fig. 10a). First, we showed that α- and β-diversities had no significant difference between the two cohorts, in which α-diversity tended to increase in the validation cohort compared with the primary cohort (Extended Data Fig. 10b,c). The α-diversity score tended to decrease in the NMDAra-treated subgroup and increase in the A+N-treated subgroup, similar to that in the primary cohort (Extended Data Fig. 10d and Fig. 1c). Moreover, a high correlation coefficient (*R*^2^) was observed for the weighted UniFrac distance between the untreated and AChEi-treated subgroups and for the unweighted UniFrac distance between the untreated and NMDAra-treated subgroups (Extended Data Fig. 10e,f). Notably, the overall microbiome composition showed a significant difference between the AChEi- and NMDAra-treated subgroups in the validation cohort, which was comparable to that in the primary cohort (weighted UniFrac distance, *R*^2^ = 0.103, *P* = 0.026; unweighted UniFrac distance, *R*^2^ = 0.057, *P* = 0.020; Extended Data Fig. 10e,f). Thus, the alteration of the salivary microbiome in anti-dementia drug-treated patients with cognitive impairment may be similar in different cohorts.

Subsequently, we explored the characteristic species in the altered salivary and gut microbiomes associated with anti-dementia drug use by multivariate analysis using confounding factors, including age, sex, disease, and cohort, in the primary cohort. We found that the abundances of *Peptostreptocccus* and *G_undefined_Clostridiales Family XIII. Incertae Sedis* significantly increased in the salivary microbiome of the AChEi-treated subgroup, whereas *Lactobacillus, Tannerella, Parvimonas, Bacteroides*, and *Bulleidia* significantly increased in the NMDAra-treated subgroup (*P* < 0.05; Fig. 3a,b). Notably, in the A+N-treated subgroup, the abundance of *G_undefined_Clostridiales Family XIII. Incertae Sedis* and *Peptostreptococcus* increased and that of *Streptococcus* decreased, as in the AChEi-treated subgroup (Fig. 3c). We also observed that the overall salivary microbiome composition was no significant difference between the AChEi- and A+N-treated subgroups (Fig. 1d,e), suggesting that the effect of AChEi use was more prominent than that of NMDAra use in patients receiving dual therapy. In the gut microbiome, *Megasphaera* was commonly enriched in the untreated subgroup, and *Faecalicatena*, *Enterococcus*, and *G_undefined*_*Lachnospiraceae* were enriched in the untreated subgroup compared with the NMDAra- and A+N-treated subgroups, whereas *G_undefined_Ruminococcaceae* and *Tidjanibacter* were increased only in the NMDAra-treated subgroup (Fig. 3d,e,f). Interestingly, most species significantly increased in patients with anti-dementia drug use in the salivary microbiome, but most of the species decreased in the gut microbiome. Thus, these findings suggest that the effects of anti-dementia drug use may differ between salivary and gut microbiome.

**Fig. 3.**
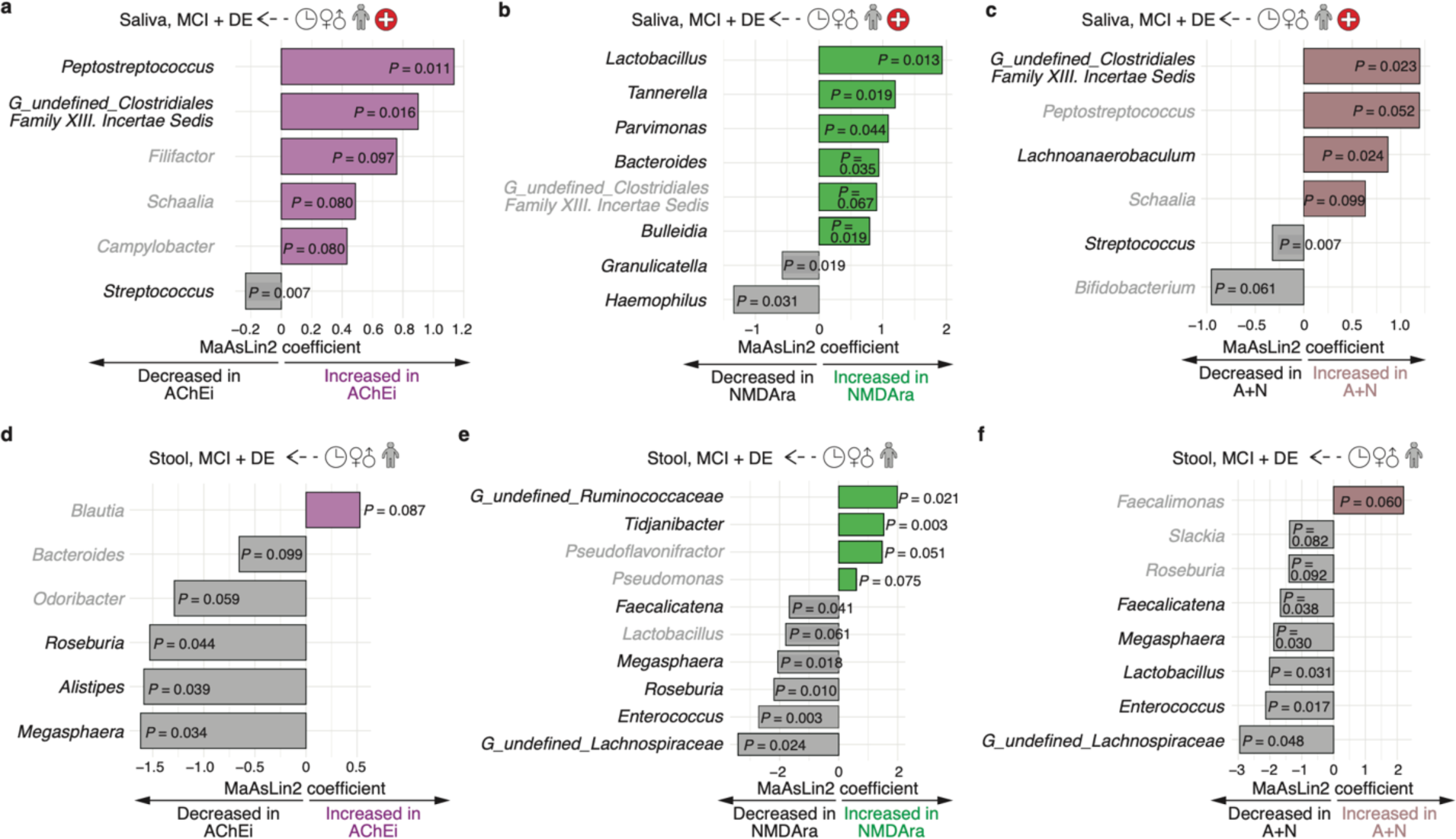
Alteration of microbial abundance depends on the type of anti-dementia drugs used. **a, b, c,** Graphs show the representative abundance of the salivary microbiome at genus level between UT vs AChEi-treated (a), UT vs NMDAra-treated (b), and UT vs A+N-treated (c) subgroups in cognitive impairment groups (MCI and DE, UT, *n* = 63; AChEi, *n* = 62; NMDAra, *n* = 27; A+N, *n* = 23, *P* < 0.1). Statistical significance was determined via the MaAsLin2 package with age, sex, disease, and cohort as random effects (*P* < 0.05), using the top 44 abundant genera with a relative mean abundance > 0.1%. **d, e, f,** Graphs show the representative abundance of the gut microbiome at genus level between UT vs AChEi-treated (d), UT vs NMDAra-treated (e), and UT vs A+N-treated subgroups (f) in cognitive impairment groups (MCI and DE, UT, *n* = 14; AChEi, *n* = 39; NMDAra, *n* = 20; A+N, *n* = 22, *P* < 0.1). Statistical significance was determined via the MaAsLin2 package with age, sex, and disease as random effects (*P* < 0.05), using the top 79 abundant genera with a relative mean abundance > 0.1%. Light characters indicate no significance (*P* > 0.05). MCI; mild cognitive impairment; DE, dementia; AChEi, acetylcholinesterase inhibitor; NMDAra, N-methyl-D-aspartate receptor antagonist; A+N, AChEi plus NMDA dual therapy; UT, untreated.

### Effect of acetylcholine and glutamate

We next focused on the salivary microbiome in patients with Parkinson’s disease because anti-parkinsonian drugs include NMDAra and anticholinergics^34^. We classified 10 anti-parkinsonian drugs based on their mode of action^34^ and defined patients with de novo Parkinson’s disease as the untreated group. Almost all patients in the treated group received levodopa and dopamine agonists. Hence we classified them into 11 drug categories based on the drug of interest (including amantadine HCl, anticholinergics, and zonisamide, an anticonvulsant), with a background of levodopa and dopamine agonists dosing (Extended Data Fig. 11a,b). We first compared the α- and β-diversities between the untreated and treated groups and found no significant changes (Extended Data Fig. 11c,d). The α-diversity score was remarkably lower in patients receiving amantadine HCl (Drug 8 subgroup) than in those receiving levodopa monotherapy (Drug 2 subgroup), which was consistent with the results for patients with cognitive impairment receiving NMDAra drugs (Extended Data Fig. 11e and Fig. 1c). Meanwhile, the β-diversity for the Drug 8 subgroup showed a high correlation coefficient, with no significance (Extended Data Fig. 11f).

Finally, we performed multivariate analyses with several confounding variables, including age, sex, disease severity, disease duration, and anti-parkinsonian drug use. The use of anticholinergics as well as age and disease duration significantly contributed to the species composition in the salivary microbiome (cumulative adjusted *R*^2^: age, 5.26%, *FDR* = 0.025; anticholinergics, 10.90%, *FDR* = 0.025; duration, 13.08%, *FDR* = 0.113; Fig. 4a). Furthermore, by analysing individual drugs as confounding factors, we observed that each drug altered the abundance of unique species (Fig. 4b). These results suggest that the drugs that modulate cerebral acetylcholine and glutamate levels affect the salivary microbiota.

**Fig. 4.**
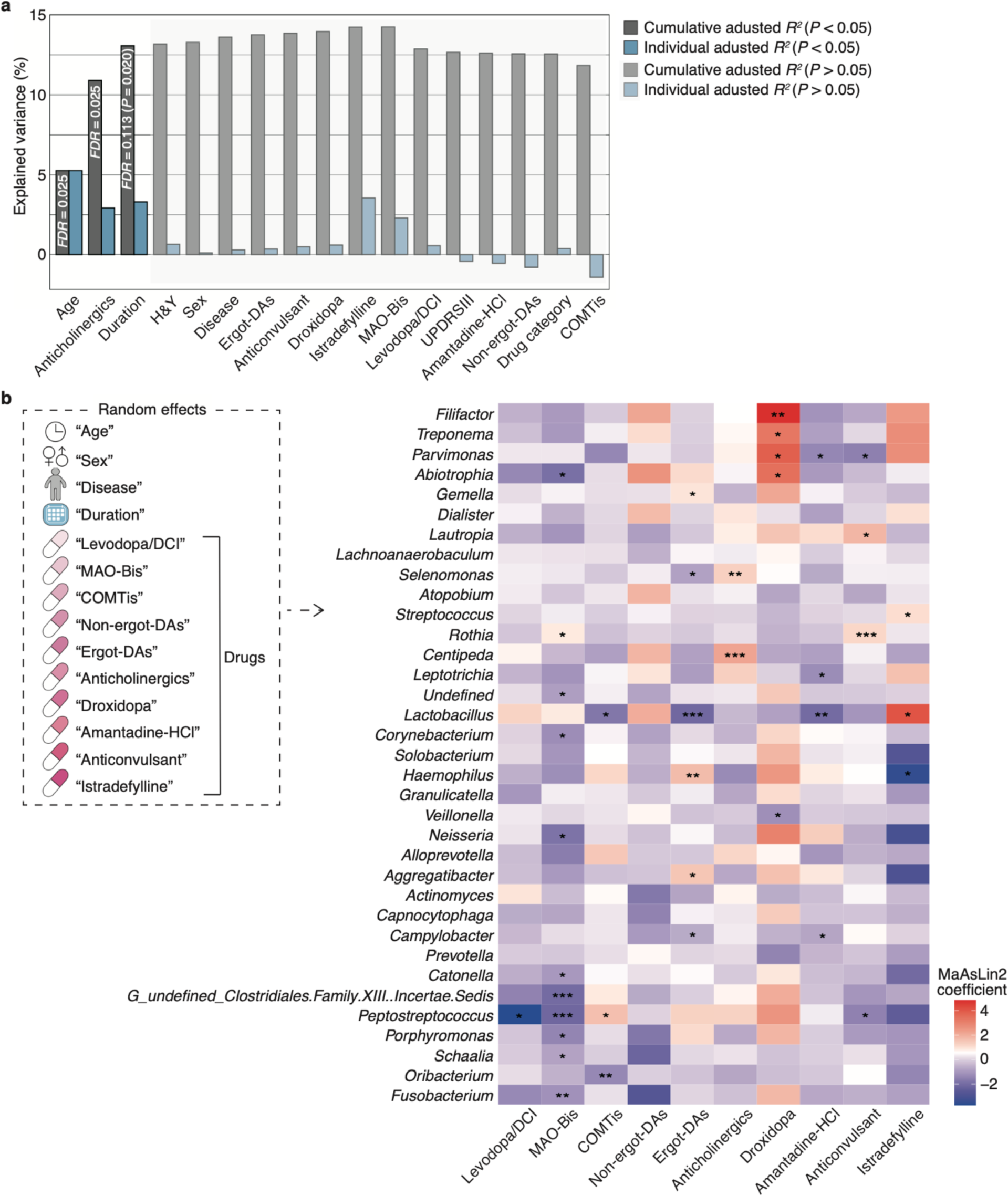
Anti-parkinsonian drug use affects salivary microbiome composition. **a,** Individual and cumulative adjusted *R*^2^ (explained variance) of covariates in stepwise RDA using the species composition of the salivary microbiome in patients with Parkinson’s disease (*n* = 62). Light colour indicates no significance (*FDR* > 0.05 and *P* > 0.05). **b,** The heatmap shows representative genera, the top 35 abundant genera with a relative mean abundance > 0.1%, enriched and depleted among patients using different anti-parkinsonian drugs (levodopa/DCI, *n* = 49; MAO-Bi, *n* = 13; COMTi, *n* = 10; non-ergot-DA, *n* = 1; ergot-DA, *n* = 33; anticholinergics, *n* = 9; droxidopa, *n* = 2; amantadine HCl, *n* = 15; anticonvulsant, *n* = 13; istradefylline, *n* = 2). Statistical significance was determined using the MaAsLin2 package with age, sex, disease type (de novo or Parkinson’s disease), disease duration, and individual drug use as random effects (*P* < 0.05). \**P* < 0.05; \*\**P* < 0.01; \*\*\**P* < 0.005. H&Y, Hoehn-Yahr scale score; DA, dopamine agonist; MAO-Bi, monoamine oxidase type B inhibitor; DCI, dopa-decarboxylase inhibitor; UPDRSⅢ, MDS Unified Parkinson’s Disease Rating Scale Part III score; COMTi, catechol-*O*-methyl transferase inhibitor.

## Discussion

The main finding of this study was that the use of anti-dementia drugs was significantly associated with human microbiome alterations and that the influence of the drugs was more prominent in the salivary microbiome than in the gut microbiome. Although large-scale meta-analyses have shown that the use of several drugs is associated with changes in the gut microbiome^30–32^, the relationship between CNS drug use and microbiome alteration remained unclear. This study may be the first report on the association of anti-dementia drug use and the human microbiome.

AChEis such as donepezil reversibly inhibit acetylcholinesterase, increasing the cerebral acetylcholine level and enhancing cholinergic neurotransmission in patients with cognitive impairment^33^. Conversely, anticholinergics normalise the hyperactivity of cholinergic neurotransmission caused by depleting dopamine in patients with Parkinson’s disease^37,38^. NMDAras, including memantine and amantadine HCl, uncompetitively block NMDA receptor activity caused by excess glutamate in patients with neurodegeneration^33,39^. These orally administered CNS drugs are absorbed in the gastrointestinal tract^33,35,36^, after which donepezil and galantamine are mainly metabolised by cytochrome P450 in the liver^40^, and NMDAras remain unmetabolised by hepatic enzymes and are excreted by the kidneys^36,41^. Based on the mode of action and metabolism of CNS drugs that regulate the brain physiology, our main finding suggested the presence of the microbiome–oral–brain axis in neurodegeneration, although it is unclear whether the mechanism for interplay between the salivary microbiome and drug use is direct or indirect. Previous studies have shown the abnormal aggregation of α-synuclein and tau in the salivary glands of patients with neurodegeneration^42,43^. Cholinergic innervation by cranial nerves Ⅶ and Ⅸ acts on the salivary glands, controlling saliva secretion^44,45^. Neurodegeneration-related proteins, such as α-synuclein, tau, and amyloid β, are secreted into the saliva in patients with neurodegenerative diseases^45^. Moreover, Schloss et al. showed that inhibiting acetylcholinesterase reduced the systemic supply of inflammatory myeloid cells in cohort studies of donepezil-treated patients with Alzheimer’s disease and in mouse experiments^46^. Collectively, as the brain controls the immune response through neurotransmitters and innervation^29,47^, the salivary microbiome may better reflect the systemic physiological changes due to the action of the CNS drugs compared with the gut microbiome. Furthermore, NMDAra-treated patients showed lowed species richness and evenness in the salivary microbiome (Fig. 1c and Extended Data Fig. 11e). Interestingly, previous studies have shown a higher α-diversity in the gut and salivary microbiome in patients with Parkinson’s diseases than in healthy controls^27,48,49^, which was consistent with our results in patients with cognitive impairment (Extended Data Fig. 3b, 7b). These results imply that the reduction in salivary microbiome diversity induced by NMDAra dosing may be correlated with therapeutic efficacy.

The main limitation of this study was that the influence of therapeutic drugs other than CNS drugs was not considered. We must evaluate the effects of various drugs on the salivary microbiome because of multimorbidity associated with their use in older individuals. Our multivariate analysis, with the use of CNS drugs as a confounding factor, highlighted that the salivary microbiome profiles are of great use for noninvasive biomarkers for monitoring the effect of drug intake and degree of cognitive function progression observed in heterogeneous and complex neurodegeneration with high accuracy.

## Methods

### Study design

Participants were enrolled in two regions: metropolitan and country regions. We defined the 23 wards of Tokyo as metropolitan areas. Patients in the metropolitan cohort were recruited at the Juntendo Tokyo Koto Geriatric Medical Center and the Memory Clinic Ochanomizu, both in Tokyo, Japan. Patients in the country cohort were recruited at the Memory Clinic Toride, Ibaraki, Japan. The study participants were adults aged 55–91 years diagnosed with MCI and dementia using cognitive function tests described below. We excluded patients with Lewy body dementia, frontotemporal dementia, and Parkinson’s disease with dementia diagnosed based on clinical data, including single photon emission computed tomography imaging and interviews from our study. Healthy controls were individuals who exhibited no cognitive impairment based on cognitive function tests and volunteers not suffering from any neurodegenerative diseases and living with the patients. The validation cohort included participants aged 41–96 years, recruited from different time periods at metropolitan hospitals and diagnosed with MCI and dementia using the same criteria in the primary cohort. Furthermore, we enrolled patients with Parkinson’s disease diagnosed using standard criteria at Juntendo University Hospital, Tokyo, Japan. The following metadata were collected for cognitive impairment assessment: age, sex, MMSE score, revised Hasegawa Dementia Scale (HDS-R) score, Alzheimer’s Disease Assessment Scale-Cognitive Subscale (ADAS-cog) score, Wechsler Adult Intelligence Scale (WAIS) score, and prescription drugs, including anti-dementia drugs. The following data were collected for Parkinson’s disease assessment: age, sex, disease duration, Hoehn-Yahr scale score, MDS Unified Parkinson’s Disease Rating Scale (MDS-UPDRS) score for Part III, and prescription drugs, including anti-parkinsonian drugs.

We divided patients with cognitive impairment into four groups according to the type of anti-dementia drug used: untreated, AChEi (donepezil, galantamine, or rivastigmine)-treated, NMDAra (memantine)-treated, and A+N-treated group. Patients with Parkinson’s disease were also divided into two groups, untreated and treated, and classified into 10 subgroups based on anti-parkinsonian drug types used in the treated group (Extended Data Fig. 11b).

### Assessment of cognitive function

The participants were divided into three groups according to MMSE and HDS-R scores: HC (MMSE ≥ 28 and HDS-R ≥ 21), MCI (24 ≤ MMSE ≤ 27 and HDS-R ≥ 21), and dementia (MMSE ≤ 23 and HDS-R ≤ 20) groups. Moreover, some participants were tested using the ADAS-cog and WAIS to confirm the robustness of the cohort.

### Sample collection and 16S rRNA amplicon sequencing

All samples were collected during the daytime, excluding those of participants who took antibiotics within 2 weeks before sampling. The saliva and stool samples were collected, frozen in liquid nitrogen, and stored at -80 ℃ until the DNA was extracted. Saliva and stool sampling, freezing, and DNA extraction based on the enzymatic lysis method from frozen samples were performed as previously described^19,50^. In brief, the V1-V2 region of the 16S rRNA gene was amplified by polymerase chain reaction using the following primer sets: forward 27Fmod (5′-AATGATACGGCGACCACCGAGATCTACACxxxxxxxxACACTCTTTCCCTACACGACGC TCTTCCGATCTagrgtttgatymtggctcag-3′) and reverse 338R (5′-CAAGCAGAAGACGGCATACGAGATxxxxxxxxGTGACTGGAGTTCAGACGTGTGCTCT TCC GATCTtgctgcctcccgtaggagt-3′), containing the Illumina Nextera adapter sequence and a unique 8-bp index sequence for each sample (indicated by xxxxxxxx). Thermal cycling was performed on a 9700 PCR system (Life Technologies, Carlsbad, CA, USA) using Ex Taq polymerase (Takara Bio, Tokyo, Japan) with the following cycling conditions: initial denaturation at 96 °C for 2 min; 25 cycles of denaturation at 96 °C for 30 s, annealing at 55° C for 45 s, and extension at 72 °C for 1 min; and final extension at 72 °C. The amplicons were purified using AMPure XP magnetic purification beads (Beckman Coulter, Brea, CA, USA) and quantified using the Quant-iT PicoGreen dsDNA Assay Kit (Life Technologies, Japan). The amplicon pools were sequenced using the Illumina Miseq Platform (2×300 bp), according to the manufacturer’s instructions.

### Data processing

We used an analysis pipeline for MiSeq barcoded amplicon sequencing of the V1-V2 region for data processing, as previously described^51,52^. Briefly, the paired-end 16S rRNA amplicon sequences were assigned to each sample based on the barcode sequence; thereafter, the reads with an average quality value < 25, mismatches to both universal primers, and possible chimeric reads were removed. Reads with BLAST match lengths of < 90% with the representative sequence in the 16S databases (RDP v10.27, CORE, and NCBI Refseq) were considered as chimaeras and removed. Finally, we adopted 10,000 filter-passed reads per sample of high-quality reads and deposited them in the DDBJ/GenBank/EMBL database. To obtain the taxonomic assignments of the OTUs, we clustered 10,000 filter-passed reads using the UCLUST algorithm with a 97% identity threshold and generated them from similarity searches against the NCBI RefSeq database downloaded on January 8, 2020, using the GLSEARCH programme.

### Microbiome and metadata analysis

We analysed microbiome diversity as previously described^50^. Briefly, UniFrac distance analysis was used to determine the dissimilarity (distance) between each pair of samples. Dissimilarity in the microbiome composition was visualised using principal coordinate analysis based on UniFrac distance analysis. Statistical significance was obtained via permutational multivariate analysis of variance, and *P*-values were adjusted using the Benjamini–Hochberg method. The OTU number was estimated using the vegan package (v2.6−4) of the statistical programming language R, version 4.0.3 (2020-10-10), based on the Chao 1 estimator.

To classify our participants based on their microbiome composition, we generated a random forest model using the randomforest package (v4.7−1.1). The AUC of the receiver-operating characteristic curves were used to find the best model to discriminate between MCI and dementia in the untreated cognitive impairment group. The best model was evaluated using the mean AUC in a 10-fold cross-validation test repeated 20 times.

Stepwise RDA was performed to evaluate the confounding variables that contributed to the microbiome composition using the ordiR2step function in the vegan package (v2.6−4), as previously described^32,53^. Distance-based RDA was performed using the Bray-Curtis distance with 999 permutations, and the *P*-values were corrected using the Benjamini–Hochberg method.

Multivariate analysis of the microbial community was performed via the R package MaAsLin2 (v1.10.0) using generated linear and mixed models^54^, as previously described^55^. The cognitive impairment model included fixed (disease [healthy control, MCI, and dementia] or drug [untreated, AChEi, NMDAra, and A+N]) and random (age, sex, cohort, disease, or drug) effects. The Parkinson’s disease model included fixed (individual drugs [levodopa/dopa-decarboxylase inhibitor, monoamine oxidase type B inhibitor, catechol-*O*-methyl transferase inhibitor, non-ergot-DA, ergot-DA, anticholinergics, droxidopa, amantadine HCl, anticonvulsant, or istradefylline]) and random (age, sex, disease [de novo Parkinson’s disease or Parkinson’s disease], disease duration, or individual drugs) effects. The random effects were illustrated in each figure.

### Statistical analysis

All statistical analyses were performed using the statistical programming language R version 4.0.3 (2020-10-10). Statistical significance was determined using Tukey’s test, Welch’s *t*-test, and Wilcoxon rank-sum test via the Benjamini–Hochberg method. The false discovery rate (*FDR*) and *P*-values were set below 0.05.

### Study approval

This study was approved by the Ethics Committee of Juntendo University School of Medicine (approval number H19-0244) and Juntendo Tokyo Koto Geriatric Medical Center (approval number 110-11). All participants provided written informed consent for study participation. The procedures in this study were performed following the principles of the Declaration of Helsinki.

### Data availability

Bacterial 16S rRNA sequencing data generated for this study were deposited in the DDBJ/GenBank/EMBL database (accession numbers: DRA016578, DRA016579, and DRA016580). Clinical data, except within this article, are not available in a public repository or supplementary material to protect the privacy and confidentiality of the study participants. Requests for clinical data can be directed to the corresponding authors and will be reviewed by the Ethics Committee of Juntendo University School of Medicine. All data shared will be de-identified.

## Acknowledgements

We would like to acknowledge all the study participants. We are grateful to the medical staff of the Department of Neurology at the Juntendo University Faculty of Medicine, Department of Gastroenterology at the Juntendo Tokyo Koto Geriatric Medical Center, and the Memory Clinic Ochanomizu. We thank Hiroaki Masuoka (Laboratory for Microbiome Sciences, RIKEN Center for Integrative Medical Sciences) for providing the script of the MaAsLin2 package for the statistical programming language R, version 4.0.3 (2020-10-10). This work was supported by the Cabinet Office of the Japanese Government, Public/Private R&D Investment Strategic Expansion PrograM: PRISM (grant number 19AC5003 to M.H. and C.A.), and was partially supported by the Japan Society for the Promotion of Science (JSPS)/The Ministry of Education, Culture, Sports, Science and Technology (MEXT), KAKENHI Grant-in-Aid for Early-Career Scientists (grant number 21K15888 to D.H.), and the Japan Agency for Medical Research and Development (AMED) (21wm0425015 to T.H. and 21dm0207070 to N.H.). We also acknowledge financial support from the JGC Japan Corporation, Japan; Otsuka Holdings Co., Ltd., Japan; and ITOCHU Chemical Frontier Corporation, Japan.

## Author contributions

D.H. and C.A. designed and conceptualised the study. D.H. performed the overall data analysis. Y.O. and W.S. conducted the 16S rRNA amplicon sequencing and data processing using an analysis pipeline. H.T.A., T.H., D.A., and T.A. performed clinical examinations and provided clinical information. Y.O. and Y.N. recruited participants. N.S., N.H., and M.H. supervised the study. D.H. wrote the manuscript. N.H., M.H., and C.A. reviewed the manuscript. All the authors have read and approved the final draft of the manuscript.

