## Extended Data Figures for "Connection between drug-mediated neurotransmission and salivary microbiome"

### Affiliations

### Extended Data Figure legends

**a**

| | 1st (Metropolis) | | | 2nd (Country) | | | Whole | | | |
|---|---|---|---|---|---|---|---|---|---|---|
| **Disease category** | **HC** | **MCI** | **DE** | **HC** | **MCI** | **DE** | **HC** | **MCI** | **DE** | ***P*-value** |
| Number of participants | 37 | 81 | 57 | 11 | 8 | 29 | 48 | 89 | 86 | – |
| Age (mean ± SD, years) | 73.4 ± 7.7 | 75.8 ± 6.4 | 77.4 ± 6.9 | 79.4 ± 5.4 | 78.0 ± 6.1 | 77.2 ± 6.8 | 74.8 ± 7.6 | 76.0 ± 6.4 | 77.3 ± 6.8 | 0.095–0.583 |
| Sex (% male) | 27.0 | 44.4 | 43.9 | 27.3 | 62.5 | 41.4 | 27.1 | 46.1 | 43.0 | 0.079–0.911 |
| MMSE (mean ± SD) | 27.8 ± 1.8 | 24.8 ± 2.0 | 18.4 ± 4.7 | 27.6 ± 1.4 | 23.4 ± 3.2 | 17.5 ± 3.9 | 27.8 ± 1.7 | 24.7 ± 2.2 | 18.1 ± 4.5 | $< 6.6\times10^{-7}$ |
| HDS-R (mean ± SD) | 27.8 ± 1.6 | 23.9 ± 3.4 | 17.0 ± 6.1 | 28.5 ± 0.8 | 23.8 ± 3.0 | 16.0 ± 4.5 | 28.0 ± 1.5 | 23.9 ± 3.4 | 16.6 ± 5.6 | $< 1.2\times10^{-5}$ |
| ADAS-cog (mean ± SD) | 5.6 ± 2.4 | 10.4 ± 3.6 | 18.4 ± 8.0 | 7.0 ± 2.0 | 11.1 ± 4.0 | 19.5 ± 6.6 | 6.0 ± 2.4 | 10.5 ± 3.7 | 18.8 ± 7.5 | $< 2.5\times10^{-4}$ |
| WAIS immediate (mean ± SD) | 11.6 ± 3.8 | 4.6 ± 2.6 | 2.5 ± 2.9 | 8.2 ± 2.8 | 4.2 ± 2.7 | 0.9 ± 1.6 | 11.0 ± 3.9 | 4.6 ± 2.6 | 1.9 ± 2.6 | $< 6.8\times10^{-5}$ |
| WAIS delayed (mean ± SD) | 11.6 ± 3.8 | 2.5 ± 2.6 | 0.7 ± 1.9 | 5.8 ± 4.1 | 1.8 ± 2.7 | 0.1 ± 0.2 | 10.5 ± 4.6 | 2.4 ± 2.7 | 0.5 ± 1.5 | < 0.006 |
| **Drug (Number of participants)** | | | | | | | | | | |
| UT | 35 | 49 | 7 | 6 | 3 | 4 | 41 | 52 | 11 | – |
| AChEi | 2 | 23 | 22 | 5 | 4 | 13 | 7 | 27 | 35 | – |
| NMDAra | 0 | 5 | 11 | 0 | 1 | 10 | 0 | 6 | 21 | – |
| A+N | 0 | 4 | 17 | 0 | 0 | 2 | 0 | 4 | 19 | – |

**b**

| | Untreated | | | |
|---|---|---|---|---|
| **Disease category** | **HC** | **MCI** | **DE** | ***P*-value** |
| Number of participants | 41 | 52 | 11 | – |
| Age (mean ± SD, years) | 73.6 ± 7.4 | 76.3 ± 5.5 | 76.6 ± 6.0 | 0.126–0.984 |
| Sex (% male) | 29.3 | 42.3 | 63.6 | 0.097–0.410 |
| MMSE (mean ± SD) | 28.0 ± 1.7 | 25.5 ± 1.3 | 19.3 ± 5.6 | $< 5.0\times10^{-6}$ |
| HDS-R (mean ± SD) | 28.0 ± 1.5 | 24.9 ± 2.9 | 18.3 ± 6.4 | < 0.001 |
| ADAS-cog (mean ± SD) | 5.4 ± 1.9 | 9.3 ± 3.3 | 15.7 ± 7.8 | $< 1.9\times10^{-4}$ |
| WAIS immediate (mean ± SD) | 11.7 ± 3.6 | 5.4 ± 3.2 | 3.1 ± 4.7 | $< 1.5\times10^{-6}$ (vs HC)<br>0.351 (MCI vs DE) |
| WAIS delayed (mean ± SD) | 11.5 ± 4.1 | 3.6 ± 3.3 | 2.1 ± 3.7 | $< 3.7\times10^{-7}$ (vs HC)<br>0.656 (MCI vs DE) |

**c**

| | MCI + DE | | | | |
|---|---|---|---|---|---|
| **Drug category** | **UT** | **AChEi** | **NMDAra** | **A+N** | ***P*-value** |
| Number of participants | 63 | 62 | 27 | 23 | – |
| Age (mean ± SD, years) | 76.3 ± 5.6 | 75.9 ± 7.8 | 78.9 ± 7.0 | 76.8 ± 5.1 | 0.210–0.986 |
| Sex (% male) | 46.0 | 50.0 | 22.2 | 52.2 | 0.073–0.998 |
| MMSE (mean ± SD) | 24.4 ± 3.5 | 21.0 ± 3.7 | 19.7 ± 4.7 | 16.5 ± 5.3 | < 0.034<br>0.551 (AChEi vs NMDAra) |
| HDS-R (mean ± SD) | 22.9 ± 5.2 | 20.9 ± 4.9 | 18.7 ± 4.9 | 13.3 ± 5.7 | < 0.016<br>0.284–0.312 (AChEi vs UT, NMDAra) |
| ADAS-cog (mean ± SD) | 11.3 ± 5.9 | 14.3 ± 7.1 | 17.8 ± 5.8 | 20.5 ± 8.0 | < 0.003<br>0.141–0.192 (AChEi vs UT, NMDAra)<br>0.524 (NMDAra vs A+N) |
| WAIS immediate (mean ± SD) | 4.6 ± 3.9 | 3.1 ± 2.4 | 1.9 ± 2.0 | 0.8 ± 1.1 | < 0.024<br>0.099–0.325 (AChEi vs UT, NMDAra)<br>0.605 (NMDAra vs A+N) |
| WAIS delayed (mean ± SD) | 3.1 ± 3.5 | 0.8 ± 1.2 | 0.6 ± 1.4 | 0.1 ± 0.2 | $< 2.1\times10^{-4}$<br>0.584–0.977 (AChEi vs NMDAra, A+N)<br>0.848 (NMDAra vs A+N) |

**Extended Data Fig. 1 | Salivary cohort demographics. a,** Upper table shows the number of participants and individual metadata values for the whole set of primary cohort. Lower table shows the number of participants in each drug category. **b,** Number of participants and individual metadata values in the UT set. **c,** Number of participants and individual metadata values in each drug category of cognitive impairment groups (MCI and DE). Statistical significance was determined using Tukey's test ($P < 0.05$). HC, healthy control; MCI, mild cognitive impairment; DE, dementia; UT, untreated; AChEi, acetylcholinesterase inhibitor; NMDAra, NMDA receptor antagonist; A+N, AChEi + NMDAra; MMSE, Mini-Mental State Examination score; HDS-R, Revised Hasegawa Dementia Scale score; ADAS-cog, Alzheimer's Disease Assessment Scale-Cognitive Subscale score; WAIS, Wechsler Adult Intelligence Scale score; SD, standard deviation.

**a**

| | Whole (Metropolis) | | | |
|---|---|---|---|---|
| **Disease category** | **HC** | **MCI** | **DE** | ***P*-value** |
| Number of participants | 28 | 31 | 64 | — |
| Age (mean ± SD, years) | 74.2 ± 9.6 | 76.4 ± 8.2 | 81.7 ± 7.3 | < 0.01 (vs DE)<br>0.56 (HC vs MCI) |
| Sex (% male) | 53.6 | 67.7 | 54.7 | 0.45–0.99 |
| MMSE (mean ± SD) | 28.6 ± 1.3 | 25.1 ± 1.7 | 17.2 ± 5.0 | < 0.003 |
| HDS-R (mean ± SD) | 27.5 ± 2.5 | 24.2 ± 3.3 | 15.3 ± 5.6 | < 0.029 |
| ADAS-cog (mean ± SD) | 6.0 ± 3.2 | 10.9 ± 3.5 | 17.9 ± 6.2 | < 0.002 |
| WAIS immediate (mean ± SD) | 9.0 ± 3.6 | 4.8 ± 2.0 | 1.7 ± 2.2 | < 0.002 |
| WAIS delayed (mean ± SD) | 7.5 ± 4.0 | 1.3 ± 1.2 | 0.5 ± 1.4 | < $1.1\times10^{-6}$ (vs HC)<br>0.60 (MCI vs DE) |
| **Drug (Number of participants)** | | | | |
| UT | 20 | 10 | 4 | — |
| AChEi | 7 | 14 | 25 | — |
| NMDAra | 1 | 4 | 16 | — |
| A+N | 0 | 3 | 19 | — |

**b**

| | Untreated | | | |
|---|---|---|---|---|
| **Disease category** | **HC** | **MCI** | **DE** | ***P*-value** |
| Number of participants | 20 | 10 | 4 | — |
| Age (mean ± SD, years) | 73.3 ± 9.8 | 73.5 ± 11.6 | 86.0 ± 4.9 | 0.080–0.998 |
| Sex (% male) | 45.0 | 80.0 | 40.0 | 0.176–0.981 |
| MMSE (mean ± SD) | 28.8 ± 1.2 | 25.9 ± 1.6 | 20.3 ± 4.8 | < 0.009 |
| HDS-R (mean ± SD) | 28.2 ± 2.2 | 24.9 ± 3.1 | 21.5 ± 5.7 | < 0.048 (vs HC)<br>0.207 (MCI vs DE) |
| ADAS-cog (mean ± SD) | 5.3 ± 2.6 | 10.7 ± 2.8 | 12.6 ± 2.7 | < $3.0\times10^{-4}$ (vs HC)<br>0.517 (MCI vs DE) |
| WAIS immediate (mean ± SD) | 10.1 ± 3.2 | 5.8 ± 2.2 | 10.5 ($n$ = 1) | 0.093 (HC vs MCI) |
| WAIS delayed (mean ± SD) | 8.6 ± 3.4 | 1.0 ± 1.1 | 6.5 ($n$ = 1) | $6.4\times10^{-4}$ (HC vs MCI) |

**c**

| | MCI + DE | | | | |
|---|---|---|---|---|---|
| **Drug category** | **UT** | **AChEi** | **NMDAra** | **A+N** | ***P*-value** |
| Number of participants | 14 | 39 | 20 | 22 | — |
| Age (mean ± SD, years) | 77.1 ± 11.6 | 79.5 ± 7.3 | 83.8 ± 6.4 | 79.2 ± 5.9 | 0.075–0.999 |
| Sex (% male) | 71.4 | 66.7 | 35.0 | 59.1 | 0.089–0.989 |
| MMSE (mean ± SD) | 24.3 ± 3.9 | 20.9 ± 4.4 | 18.8 ± 5.4 | 15.8 ± 5.9 | < 0.011 (UT vs NMDAra, A+N)<br>0.001 (AChEi vs A+N)<br>0.152–0.390 |
| HDS-R (mean ± SD) | 23.9 ± 4.3 | 19.3 ± 6.2 | 17.7 ± 5.5 | 13.1 ± 5.0 | < 0.049 (vs UT)<br>$6.0\times10^{-4}$ (AChEi vs A+N)<br>0.057–0.744 |
| ADAS-cog (mean ± SD) | 11.2 ± 2.9 | 14.9 ± 6.1 | 15.1 ± 4.2 | 20.0 ± 7.4 | < 0.012 (A+N vs UT, AChEi)<br>0.057–0.999 |
| WAIS immediate (mean ± SD) | 7.0 ± 2.8 | 3.3 ± 1.9 | 1.3 ± 1.1 | 0.9 ± 0.7 | < 0.013<br>0.059–0.975 (NMDAra vs AChEi, A+N) |
| WAIS delayed (mean ± SD) | 2.4 ± 2.6 | 1.0 ± 1.2 | 0.2 ± 0.3 | 0.1 ± 0.3 | < 0.035 (UT vs NMDAra, A+N)<br>0.236–0.999 |

**Extended Data Fig. 2 | Stool cohort demographics. a,** Upper table shows the number of participants and individual metadata values for the whole set. Lower table shows the number of participants in each drug category. **b,** Number of participants and individual metadata values in the UT set. **c,** Number of participants and individual metadata values in each drug category of cognitive impairment groups (MCI and DE). Statistical significance was determined using Tukey's test ($P < 0.05$). HC, healthy control; MCI, mild cognitive impairment; DE, dementia; UT, untreated; AChEi, acetylcholinesterase inhibitor; NMDAra, NMDA receptor antagonist; A+N, AChEi + NMDAra; MMSE, Mini-Mental State Examination score; HDS-R, Revised Hasegawa Dementia Scale score; ADAS-cog, Alzheimer's Disease Assessment Scale-Cognitive subscale score; WAIS, Wechsler Adult Intelligence Scale score; SD, standard deviation.

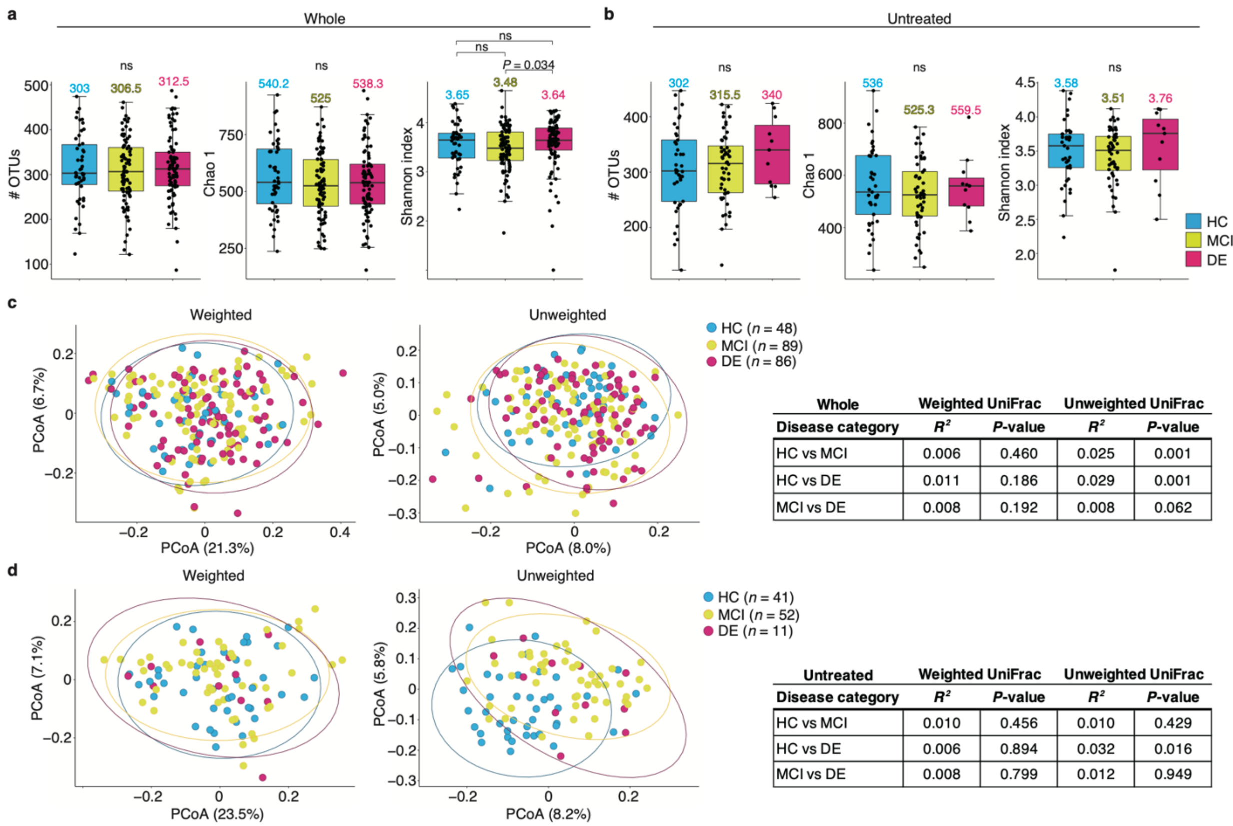


| Whole | Weighted UniFrac | | Unweighted UniFrac | |
|---|---|---|---|---|
| Disease category | $R^2$ | *P*-value | $R^2$ | *P*-value |
| HC vs MCI | 0.006 | 0.460 | 0.025 | 0.001 |
| HC vs DE | 0.011 | 0.186 | 0.029 | 0.001 |
| MCI vs DE | 0.008 | 0.192 | 0.008 | 0.062 |

| Untreated | Weighted UniFrac | | Unweighted UniFrac | |
|---|---|---|---|---|
| Disease category | $R^2$ | *P*-value | $R^2$ | *P*-value |
| HC vs MCI | 0.010 | 0.456 | 0.010 | 0.429 |
| HC vs DE | 0.006 | 0.894 | 0.032 | 0.016 |
| MCI vs DE | 0.008 | 0.799 | 0.012 | 0.949 |

**Extended Data Fig. 3 | Comparison of the salivary microbiome diversity among the disease categories. a, b,** Observed OTU number and α-diversity scores (Chao 1 and Shannon index) by disease category in the whole (a) and untreated (b) sets (whole: HC, *n* = 48; MCI, *n* = 89; DE, *n* = 86; untreated: HC, *n* = 41; MCI, *n* = 52; DE, *n* = 11). The median values are shown. Statistical significance was determined using the Wilcoxon rank-sum test with Benjamini–Hochberg correction ($P < 0.05$). ns, not significant. **c, d,** Weighted and unweighted UniFrac-PCoA between each disease category in the whole (c) and untreated sets (d). The number of participants in each disease category is shown. The $R^2$ and *P*-values were determined using permutational multivariate analysis of variance via the Benjamini–Hochberg method. Dots represent individual participants. OTU, operational taxonomic unit; HC, healthy control; MCI, mild cognitive impairment; DE, dementia; PCoA, principal coordinate analysis.

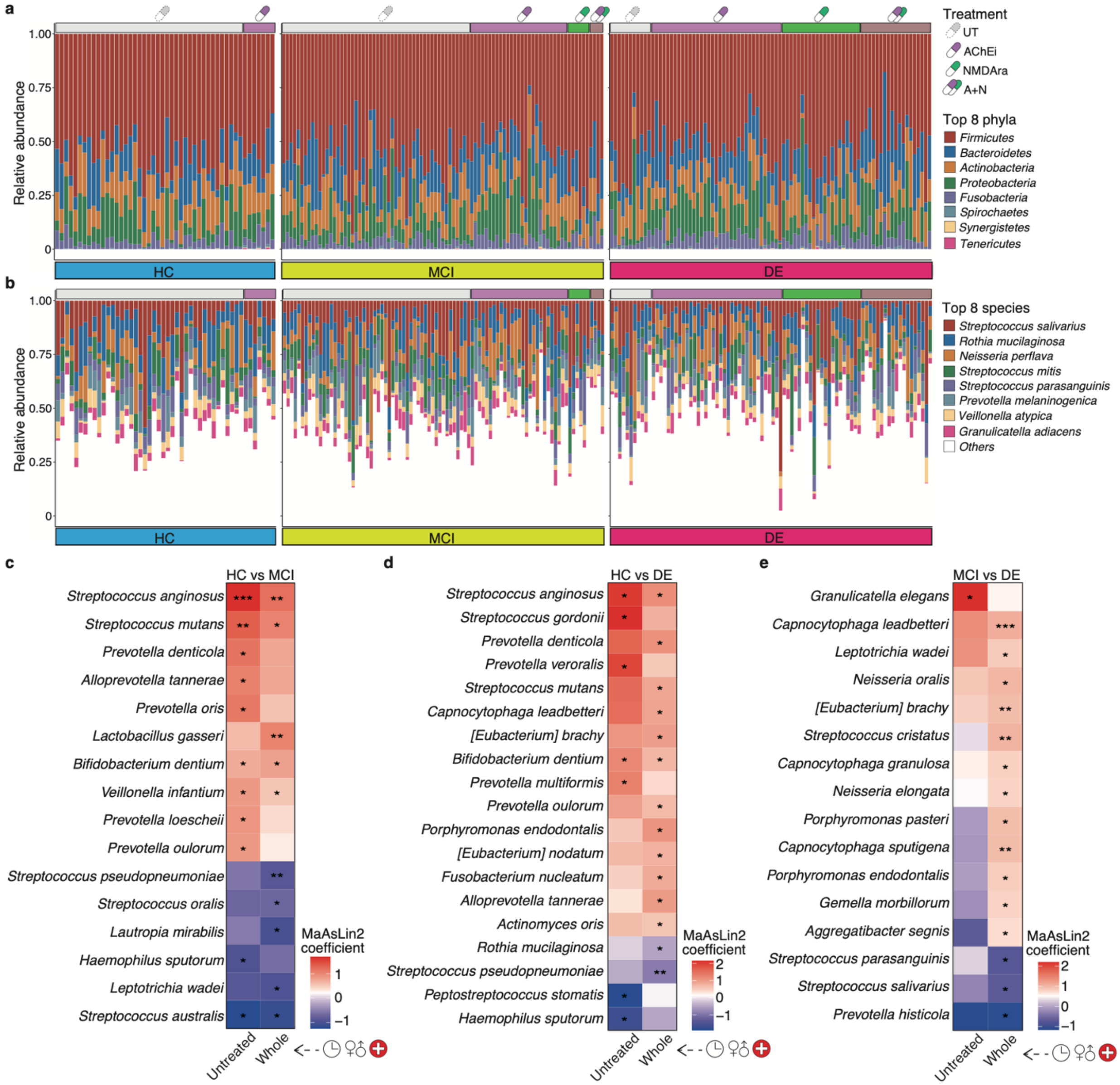


**Extended Data Fig. 4 | Salivary microbiome composition at phylum and species levels. a, b,** Relative abundance plots of salivary microbial phyla (a) and species (b) across disease categories with drug category information (HC, $n$ = 48; MCI, $n$ = 89; DE, $n$ = 86). Bars represent individual participants and their colours show the top eight most abundant phyla and the top eight most abundant species of the 101 species, with a relative mean abundance of 0.1%. **c, d, e,** Heatmaps showing representative species significantly enriched and depleted between HC vs. MCI (c), HC vs. DE (d), and MCI vs. DE groups (e) in whole sets, which contained a mixture of treated and UT patients, and untreated sets (whole: HC, $n = 48$; MCI, $n = 89$; DE, $n = 86$; UT: HC, $n = 41$; MCI, $n = 52$; DE, $n = 11$). Statistical significance was determined using the MaAsLin2 package, with age, sex, and cohort as random effects ($P < 0.05$). HC, healthy control; MCI, mild cognitive impairment; DE, dementia UT, untreated; AChEi, acetylcholinesterase inhibitor; NMDAra, NMDA receptor antagonist; A+N, AChEi + NMDAra.

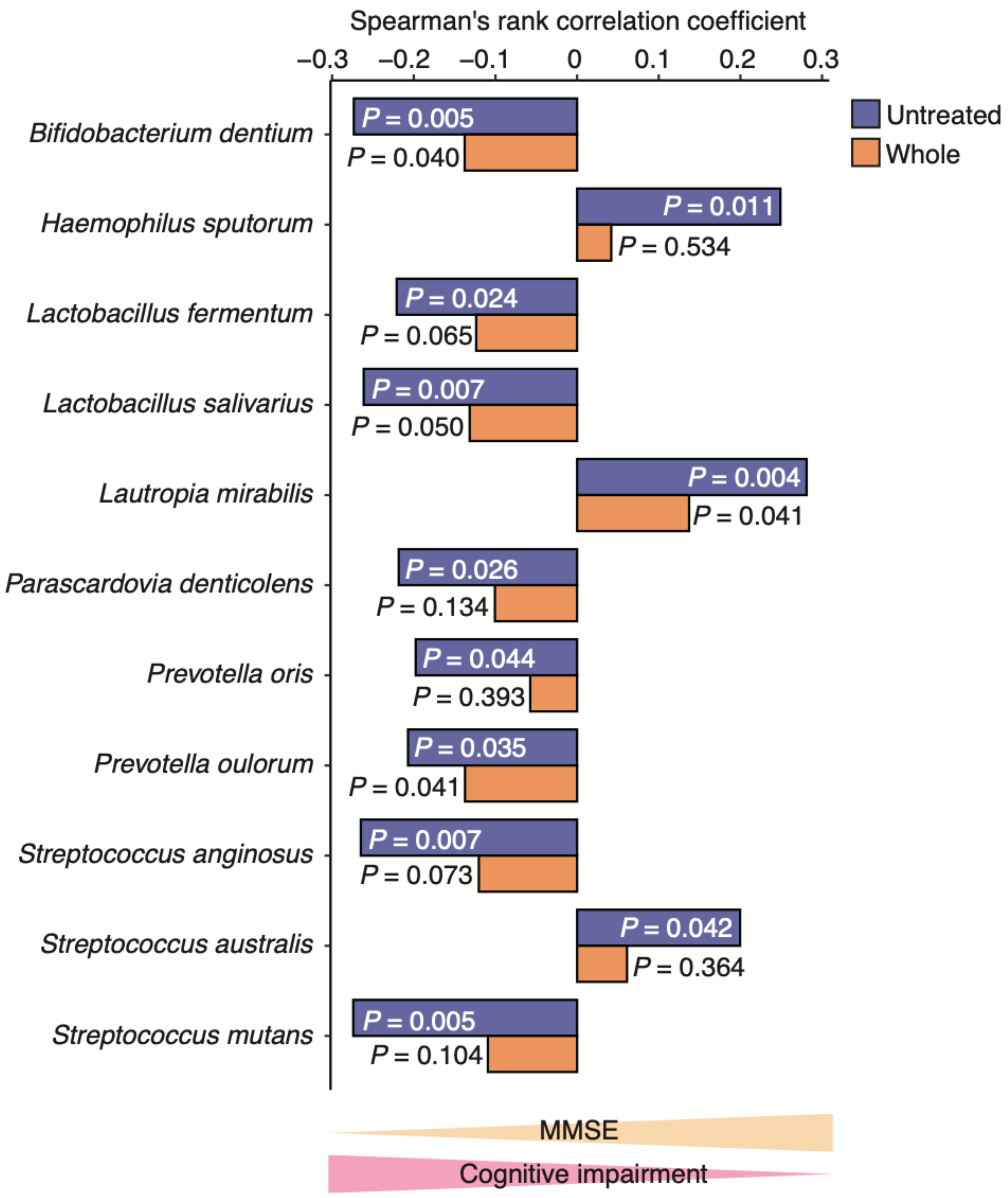


**Extended Data Fig. 5 | Correlation analysis of the salivary microbial species abundance and MMSE score.** The graph shows the representative species based on Spearman's correlation coefficients in the whole and untreated sets (whole, $n = 223$; untreated, $n = 104$). The representative species depicted in the figure were significantly different from those in the untreated set ($P < 0.05$). Lower MMSE scores indicate cognitive decline. MMSE, Mini-Mental State Examination.

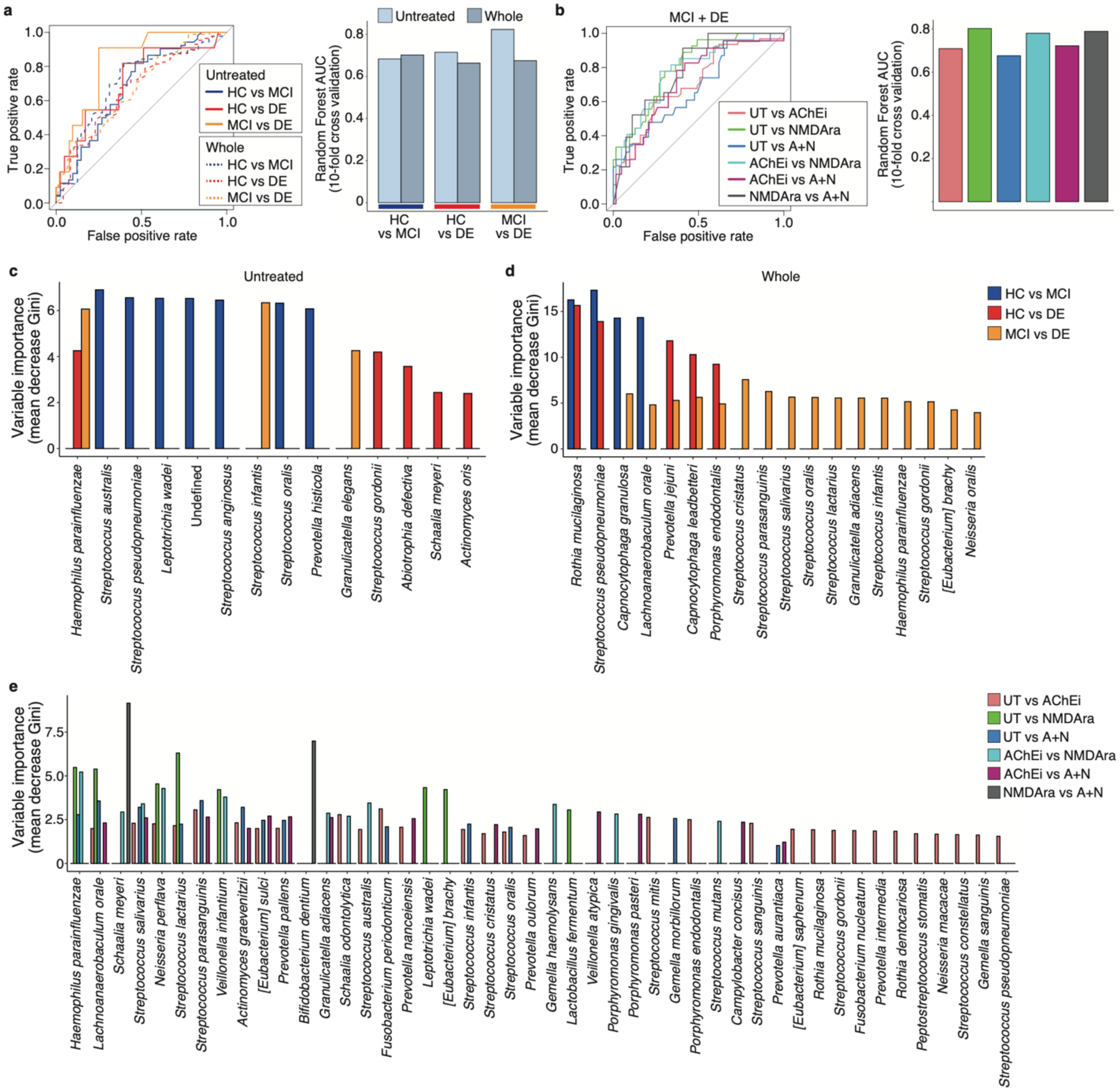


**Extended Data Fig. 6 | Discrimination of the species composition using a random forest classifier. a,** Left panel shows the receiver-operating characteristic (ROC) curve for classifying each disease category based on 20 rounds of 10-fold cross-validation in the whole and UT sets (whole: HC, *n* = 48; MCI, *n* = 89; DE, *n* = 86; UT: HC, *n* = 41; MCI, *n* = 52; DE, *n* = 11). The graph on the right shows the area under the ROC curve (AUROC). Random forest analysis was performed using the top 102 species to distinguish between the MCI and DE groups, with high performance in the UT set. **b,** Left panel shows the ROC curve for classifying each drug category based on 20 rounds of 10-fold cross-validation in the cognitive impairment groups (MCI and DE; UT, $n$ = 63; AChEi, $n$ = 62; NMDAra, $n$ = 27; A+N, $n$ = 23). The graph on the right shows the AUROC. Random forest analysis was performed using the top 102 species to distinguish between the MCI and DE groups, with high performance in the UT set, which is similar to the illustration in Extended Data Fig. 6a. **c, d, e,** Graphs show the variable importance (mean decrease Gini) of the random forest at the time of UT (c) and whole sets (d) to discriminate the disease category and each drug category (e). HC, healthy control; MCI, mild cognitive impairment; DE, dementia UT, untreated; AChEi, acetylcholinesterase inhibitor; NMDAra, NMDA receptor antagonist; A+N, AChEi + NMDAra.

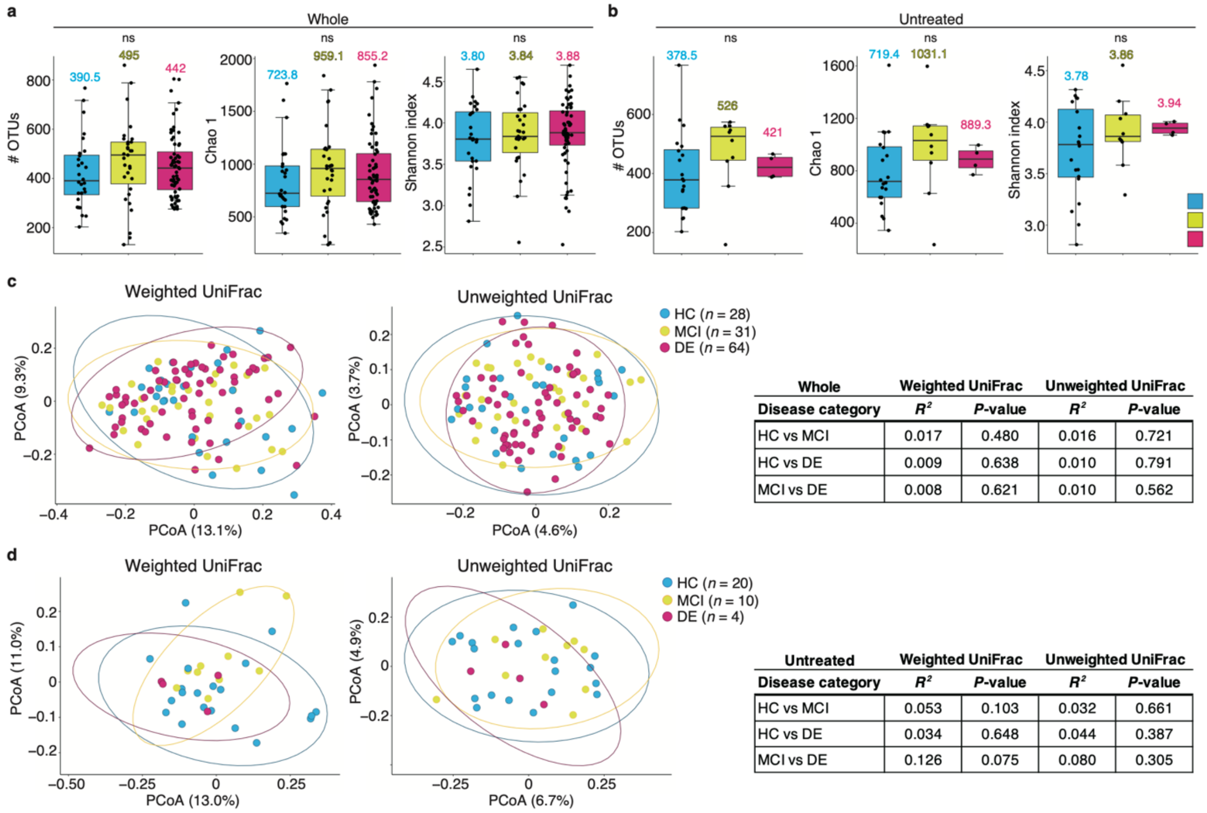


| Whole | Weighted UniFrac | | Unweighted UniFrac | |
|---|---|---|---|---|
| Disease category | $R^2$ | *P*-value | $R^2$ | *P*-value |
| HC vs MCI | 0.017 | 0.480 | 0.016 | 0.721 |
| HC vs DE | 0.009 | 0.638 | 0.010 | 0.791 |
| MCI vs DE | 0.008 | 0.621 | 0.010 | 0.562 |

| Untreated | Weighted UniFrac | | Unweighted UniFrac | |
|---|---|---|---|---|
| Disease category | $R^2$ | *P*-value | $R^2$ | *P*-value |
| HC vs MCI | 0.053 | 0.103 | 0.032 | 0.661 |
| HC vs DE | 0.034 | 0.648 | 0.044 | 0.387 |
| MCI vs DE | 0.126 | 0.075 | 0.080 | 0.305 |

**Extended Data Fig. 7 | Comparison of the gut microbiome diversity among the disease categories. a, b,** Graphs show the observed OTU number and α-diversity scores (Chao 1 and Shannon index) by disease category in the whole (a) and untreated (b) sets (whole: HC, $n = 28$; MCI, $n = 31$; DE, $n = 64$; untreated: HC, $n = 20$; MCI, $n = 10$; DE, $n = 4$). The graph shows the median values. Statistical significance was determined using the Wilcoxon rank-sum test with Benjamini–Hochberg correction ($P < 0.05$). ns, not significant. **c, d,** Weighted and unweighted UniFrac-PCoA between each disease category in the whole (c) and untreated sets (d). The number of participants in each disease category is shown. The $R^2$ and *P*-values were determined using permutational multivariate analysis of variance with Benjamini–Hochberg correction. The dots represent individual participants. HC, healthy control; MCI, mild cognitive impairment; DE, dementia; OTU, operational taxonomic unit; PCoA, principal coordinate analysis.

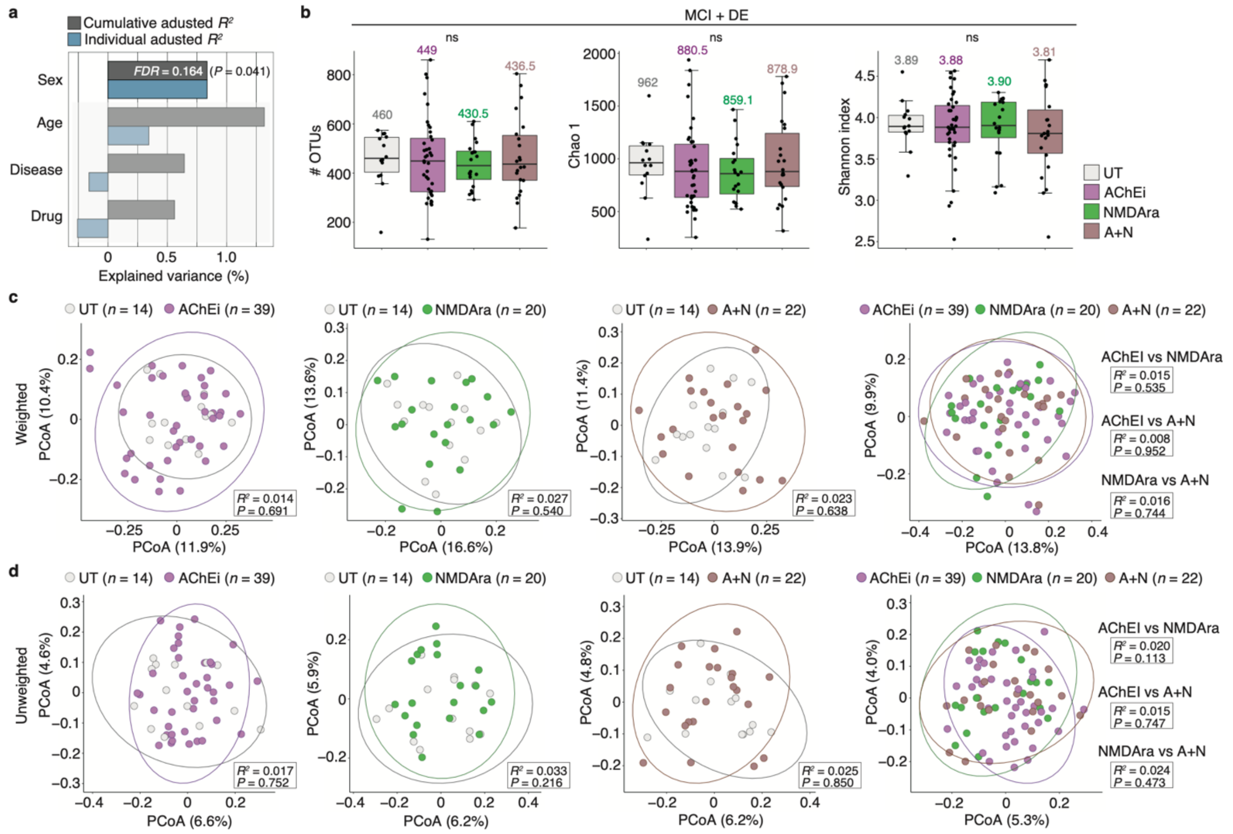


**Extended Data Fig. 8 | Contribution of anti-dementia drug use on the gut microbiome. a,** Individual and cumulative adjusted $R^2$ (explained variance) of covariates in stepwise RDA using weighted UniFrac distance in the cognitive impairment groups (MCI and DE; $n = 95$). Light colour indicates no significance ($FDR > 0.05$, $P > 0.05$). **b,** Comparison of observed OTU number and α-diversity score (Chao 1 and Shannon index) by drug category in the cognitive impairment group (MCI and DE; $n = 95$). The graph shows the median values. Statistical significance was determined using the Wilcoxon rank-sum test with Benjamini–Hochberg correction ($P < 0.05$). ns, not significant. **c, d,** Weighted (c) and unweighted (d) UniFrac-PCoA between each drug category in the cognitive impairment group (MCI and DE; $n = 95$). The number of participants in each drug category is shown. The $R^2$ and $P$-values were determined using permutational multivariate analysis of variance via the Benjamini–Hochberg method. The dots represent individual participants. HC, healthy control; MCI, mild cognitive impairment; DE, dementia; OTU, operational taxonomic unit; PCoA, principal coordinate analysis; RDA, redundancy analysis.

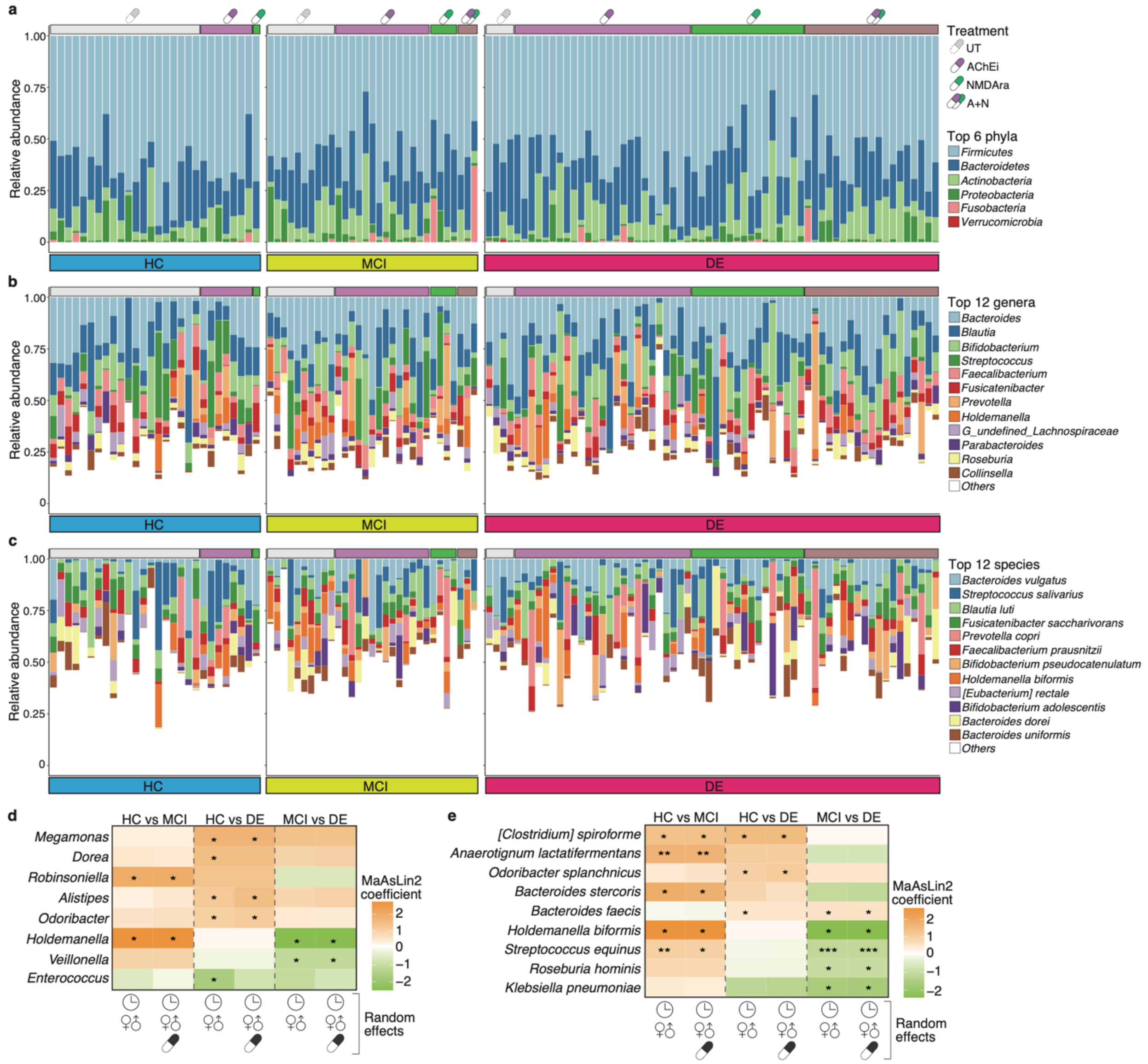


**Extended Data Fig. 9 | Gut microbiome composition of phylum, genus, and species levels. a, b, c,** Relative abundance plots of gut microbial phyla (a), genera (b), and species (c) across disease categories with drug category information (HC, *n* = 28; MCI, *n* = 31; DE, *n* = 64). Bars represent individual participants, and their colour shows the top six most abundant phyla and the top 12 most abundant bacteria of the 78 genera and 151 species, with a relative mean abundance > 0.1%. **d, e,** Heatmaps show representative genera (d) and species (e) significantly enriched and depleted in each disease category in the whole set (HC, *n* = 28; MCI, *n* = 31; DE, *n* = 64). Statistical significance was determined using the MaAsLin2 package, with age, sex, and/or drug as random effects ($P < 0.05$). *$P < 0.05$; **$P < 0.01$; ***$P < 0.005$. HC, healthy control; MCI, mild cognitive impairment; DE, dementia UT, untreated; AChEi, acetylcholinesterase inhibitor; NMDAra, NMDA receptor antagonist; A+N, AChEi + NMDAra.

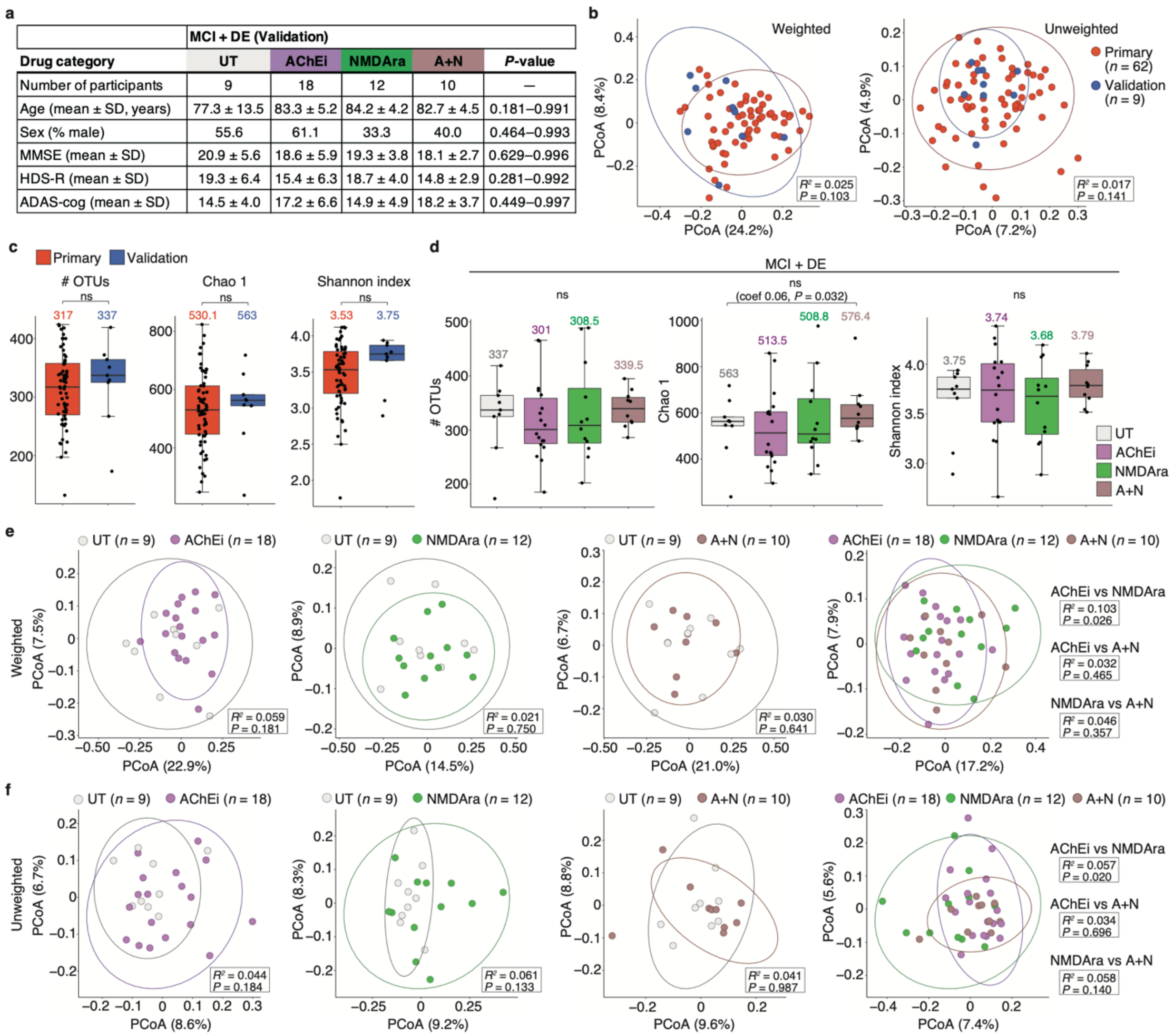


**Extended Data Fig. 10 | Contribution of anti-dementia drug use on the salivary microbiome diversity in the validation cohort. a,** Table shows the validation cohort demography, including the number of participants and individual metadata values, in the cognitive impairment groups (MCI and DE, $n$ = 49). **b,** Weighted and unweighted UniFrac-PCoA between HCs of the primary and validation cohorts (primary, $n$ = 62; validation, $n$ = 9). The $R^2$ and $P$-values were determined using permutational multivariate analysis of variance via the Benjamini–Hochberg method. **c,** Comparison of the observed OTU number and α-diversity scores (Chao 1 and Shannon index) between HCs of the primary and validation cohorts (primary, $n$ = 62; validation, $n$ = 9). **d,**

Comparison of the observed OTU number and α-diversity scores (Chao 1 and Shannon index) by drug category in the cognitive impairment groups (MCI and DE, $n = 49$). The graph shows the median values. Statistical significance was determined using the Wilcoxon rank-sum test with Benjamini–Hochberg correction and the MaAsLin2 package with age, sex, and disease as random effects ($P < 0.05$). ns, not significant. **e, f,** Weighted (e) and unweighted (f) UniFrac-PCoA between each drug category in the cognitive impairment group (MCI and DE; $n = 49$). The number of participants in each drug category is shown. The $R^2$ and $P$-values were determined using permutational multivariate analysis of variance via the Benjamini–Hochberg method. The dots represent individual participants. HC, healthy control; MCI, mild cognitive impairment; DE, dementia UT, untreated; AChEi, acetylcholinesterase inhibitor; NMDAra, NMDA receptor antagonist; A+N, AChEi + NMDAra, OTU, operational taxonomic unit; PCoA, principal coordinate analysis.

**a**

| Disease category | *de novo* (UT) | PD (Treated) | *P*-value |
|---|---|---|---|
| Number of participants | 10 | 52 | — |
| Age (mean ± SD, years) | 57.1 ± 13.5 | 64.0 ± 12.1 | 0.175 |
| Sex (% male) | 70.0 | 46.2 | 0. 179 |
| Duration (mean ± SD, years) | 1.9 ± 1.7 | 9.5 ± 5.8 | $8.84\times10^{-10}$ |
| H&Y stage (mean ± SD) | 1.6 ± 0.5 | 2.0 ± 0.7 | 0.066 |
| UPDRSIII (mean ± SD) | 23.5 ± 6.8 | 16.2 ± 8.1 | 0.012 |

**b**

| Drug category | Medication | No of participants |
|---|---|---|
| UT | - | 10 |
| Drug 2 | Levodopa/DCI | 6 |
| Drug 3 | DAs + others | 2 |
| Drug 4 | Levodopa/DCI + DAs | 6 |
| Drug 5 | Levodopa/DCI + MAO-Bis | 1 |
| Drug 6 | Levodopa/DCI + MAO-Bis + COMTis | 1 |
| Drug 7 | Levodopa/DCI + Anticholinergics + others | 6 |
| Drug 8 | Levodopa/DCI + Amantadine-HCl + others | 11 |
| Drug 9 | Levodopa/DCI + Antichorinergics + Amantadine-HCl + others | 3 |
| Drug 10 | Levodopa/DCI + Anticonvulsant + others (w/o Antichorinergics, Amantadine-HCl) | 8 |
| Drug 11 | Levodopa/DCI + MAO-Bis + COMTis + DAs + others (w/o Antichorinergics, Amantadine-HCl, Anticonvulsant) | 8 |

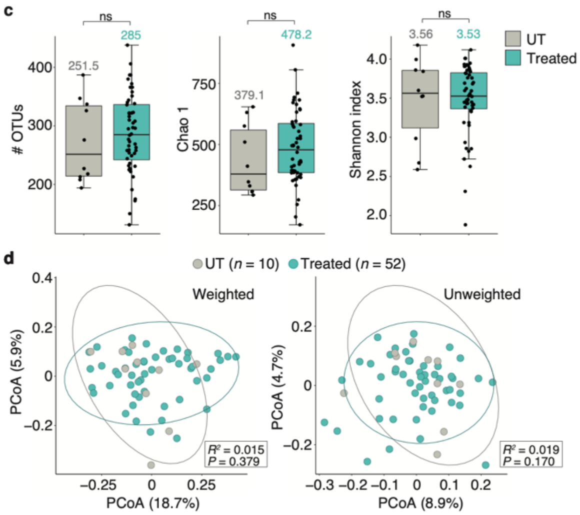


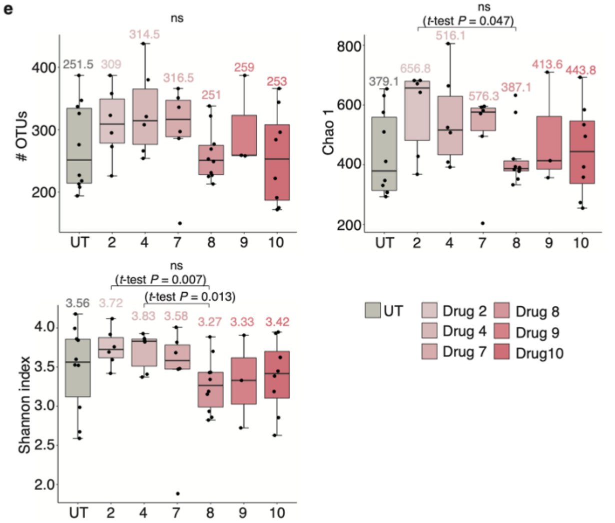


**f**

| Drug category | | Weighted UniFrac $R^2$ | Weighted UniFrac *P*-value | Unweighted UniFrac $R^2$ | Unweighted UniFrac *P*-value |
|---|---|---|---|---|---|
| UT | vs Drug 2 | 0.022 | 0.954 | 0.065 | 0.540 |
| | vs Drug 4 | 0.055 | 0.501 | 0.075 | 0.237 |
| | vs Drug 7 | 0.014 | 0.985 | 0.067 | 0.446 |
| | vs Drug 8 | 0.050 | 0.364 | 0.055 | 0.266 |
| | vs Drug 9 | 0.100 | 0.274 | 0.070 | 0.740 |
| | vs Drug 10 | 0.047 | 0.527 | 0.060 | 0.385 |
| Drug 2 | vs Drug 4 | 0.022 | 0.956 | 0.090 | 0.464 |
| | vs Drug 7 | 0.034 | 0.821 | 0.074 | 0.907 |
| | vs Drug 8 | 0.069 | 0.292 | 0.068 | 0.269 |
| | vs Drug 9 | 0.147 | 0.312 | 0.100 | 0.884 |
| | vs Drug 10 | 0.075 | 0.390 | 0.067 | 0.729 |
| Drug 4 | vs Drug 7 | 0.061 | 0.574 | 0.071 | 0.937 |
| | vs Drug 8 | 0.054 | 0.391 | 0.080 | 0.096 |
| | vs Drug 9 | 0.161 | 0.248 | 0.124 | 0.450 |
| | vs Drug 10 | 0.098 | 0.274 | 0.089 | 0.184 |
| Drug 7 | vs Drug 8 | 0.034 | 0.610 | 0.072 | 0.181 |
| | vs Drug 9 | 0.069 | 0.634 | 0.104 | 0.814 |
| | vs Drug 10 | 0.023 | 0.872 | 0.069 | 0.616 |
| Drug 8 | vs Drug 9 | 0.035 | 0.639 | 0.075 | 0.444 |
| | vs Drug 10 | 0.031 | 0.679 | 0.042 | 0.923 |

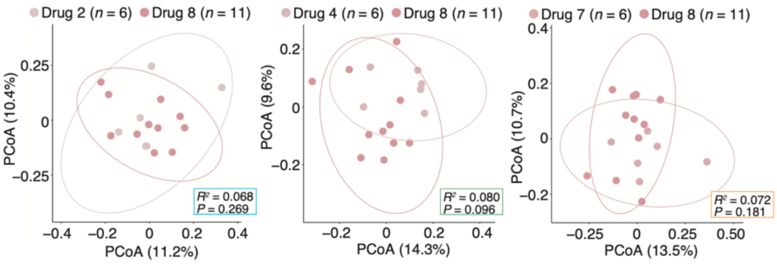


**Extended Data Fig. 11 | Contribution of anti-parkinsonian drug use on the salivary microbiome diversity in patients with Parkinson's disease. a, b,** Table shows the Parkinson's disease cohort demographics, including the number of participants and individual metadata values, in the disease (a) and drug (b) categories (*n* = 62). The de novo Parkinson's disease group was defined as the untreated group (a). Almost all patients in the treated group received levodopa and DAs, and thus we classified them into 11 subgroups based on the drug of interest (b). **c,** Comparison of the observed OTU number and α-diversity scores (Chao 1 and Shannon index)

between the UT and treated groups (UT, $n$ = 10; treated $n$ = 52). The graph shows the median values. **d,** Weighted and unweighted UniFrac-PCoA between the UT and treated groups. The numbers of participants are shown. **e,** Comparison of the observed OTU number and α-diversity scores (Chao 1 and Shannon index) by drug category of interest. The graph shows the median values. **f,** The left table shows each $R^2$ and $P$-value determined using permutational multivariate analysis of variance via the Benjamini–Hochberg method for the drug categories of interest. The right panels show representative unweighted UniFrac-PCoA. The number of participants in each drug category is shown in Extended Data Fig. 11b. To calculate the α-diversity score, statistical significance was determined using the Wilcoxon rank-sum test with Benjamini–Hochberg correction ($P < 0.05$). ns, not significant. The dots represent individual participants. UT, untreated; PD, Parkinson's disease; DA, dopamine agonist; MAO-Bi, monoamine oxidase type B inhibitor; DCI, dopa-decarboxylase inhibitor; UPDRSIII, MDS Unified Parkinson's Disease Rating Scale Part III score; COMTi, catechol-*O*-methyl transferase inhibitor.
